# Dynamics of lineage commitment revealed by single-cell transcriptomics of differentiating embryonic stem cells

**DOI:** 10.1101/068288

**Authors:** Stefan Semrau, Johanna Goldmann, Magali Soumillon, Tarjei S. Mikkelsen, Rudolf Jaenisch, Alexander van Oudenaarden

## Abstract

Gene expression heterogeneity in the pluripotent state of mouse embryonic stem cells (mESCs) has been increasingly well-characterized. In contrast, exit from pluripotency and lineage commitment have not been studied systematically at the single-cell level. Here we measured the gene expression dynamics of retinoic acid driven mESC differentiation using an unbiased single-cell transcriptomics approach. We found that the exit from pluripotency marks the start of a lineage bifurcation as well as a transient phase of susceptibility to lineage specifying signals. Our study revealed several transcriptional signatures of this phase, including a sharp increase of gene expression variability. Importantly, we observed a handover between two classes of transcription factors. The early-expressed class has potential roles in lineage biasing, the late-expressed class in lineage commitment. In summary, we provide a comprehensive analysis of lineage commitment at the single cell level, a potential stepping stone to improved lineage control through timing of differentiation cues.

## INTRODUCTION

*In vitro* differentiation is a key technology to enable the use of embryonic and induced pluripotent stem cells as disease models and for therapeutic applications ^1 2^. Existing directed differentiation protocols, which have been gleaned from *in vivo* development, are laborious and produce heterogeneous cell populations ^3^. Protocol optimization typically requires costly and time consuming trial-and-error experiments. To be able to design more efficient and specific differentiation regimens in a systematic way we require a better understanding of the decision-making process that underlies the generation of cell type diversity ^4^.

Lineage decision-making is fundamentally a single-cell process ^5^ and the response to lineage specifying signals depends on the state of the individual cell. A substantial body of work has revealed lineage biases related to, for example, cell cycle phase or pre-existing subpopulations in the pluripotent state ^6 7 8 4^. The commitment of pluripotent cells to a particular lineage, on the other hand, has not yet been studied systematically at the single-cell level. We define a cell to be committed, if its state cannot be reverted by removal of the lineage specifying signal. Here we set out to characterize the single-cell gene expression dynamics of differentiation, from exit from pluripotency to lineage commitment.

## RESULTS

As a well-characterized model system to study *in vitro* differentiation we used mouse embryonic stem cells (mESCs). To drive cells quickly out of the pluripotent state, we used all-trans retinoic acid (RA) as differentiation agent. E14 mESCs were grown feeder free in 2i medium ^9^ for several passages to minimize heterogeneity before differentiation in the basal medium (N2B27 medium) and RA (Fig. 1a). Within only 96 h the cells underwent a profound change in morphology from tight, round, homogeneous colonies to strongly adherent, morphologically heterogeneous cells (Fig. 1a). To characterize the differentiation process at the population level we measured gene expression by bulk RNA-seq at four early time points (0, 6, 12 and 24 h) as well as 6 additional time points during 96 h of continuous RA exposure (Supplementary Fig. 1a and b). Starting at 12 h the expression of most pluripotency markers declined, indicating the exit from pluripotency at that time. This was supported phenotypically by changes in morphology and cell cycle phase lengths (Supplementary Fig. 1c-e). Replating cells at clonal density in 2i medium showed that 90% of the cells exited from pluripotency between 12 h and 36 h of RA exposure (Supplementary Fig. 1f). Following exit from pluripotency, established ectoderm and extraembryonic endoderm (XEN) markers were up-regulated (Supplementary Fig. 1b), as to be expected from previous results on RA driven differentiation ^10 11^. Up-regulation of ectoderm markers started after 24 h of RA exposure, XEN markers were up-regulated only after 48 h. The more than 24 h delay between the exit from pluripotency and the expression of XEN markers suggested that a sub-population of cells went through a transitory phase between pluripotency and lineage commitment. We hypothesized that in this transient phase lineage decisions could be influenced by changes in signaling input.

To investigate this transition process in individual cells we used the recently developed Single Cell RNA Barcoding and Sequencing method ^12^ (SCRB-seq). We quantified the transcriptional profiles of over 2,000 single cells, sampled at 9 time points during differentiation, typically spaced 12 h apart. We used t-distributed stochastic neighbor embedding (t-SNE) to place individual cells with similar expression profiles in proximity to one another (Fig. 1b, Supplementary Fig 2f). To assess the pluripotency status of individual cells we used the expression level of the established pluripotency marker *rex1* ^13^. t-SNE showed that gene expression changed homogeneously throughout the population for the first 12 h of RA exposure, while *rex1* expression was overall high. Subsequently, the cells began to diverge in their gene expression profiles. Quantification of single-cell variability of gene expression confirmed a steep increase in variability between 12 h and 24 h (Fig. 1c). Coincidentally, *rex1* expression declined in the majority of cells after 12 h, indicating the cells’ exit from pluripotency. Subsequently, gene expression variability continued to increase until the end of the experiment. The t-SNE map further showed that expression profiles started to bifurcate after 24 h of RA exposure and by 96 h two separate subpopulations could be discerned. We partitioned cells into clusters using k-means clustering and confirmed by stability analysis ^14^ that there were two robust clusters. Single-cell gene expression variability within those two clusters at 96 hours was comparable to the pluripotent state (Fig. 1c). Gene expression in the two clusters was thus regulated as tightly as in the pluripotent state. Hence, we interpret the two observed clusters as two different cell types that emerged during RA differentiation.

**Figure 1.**
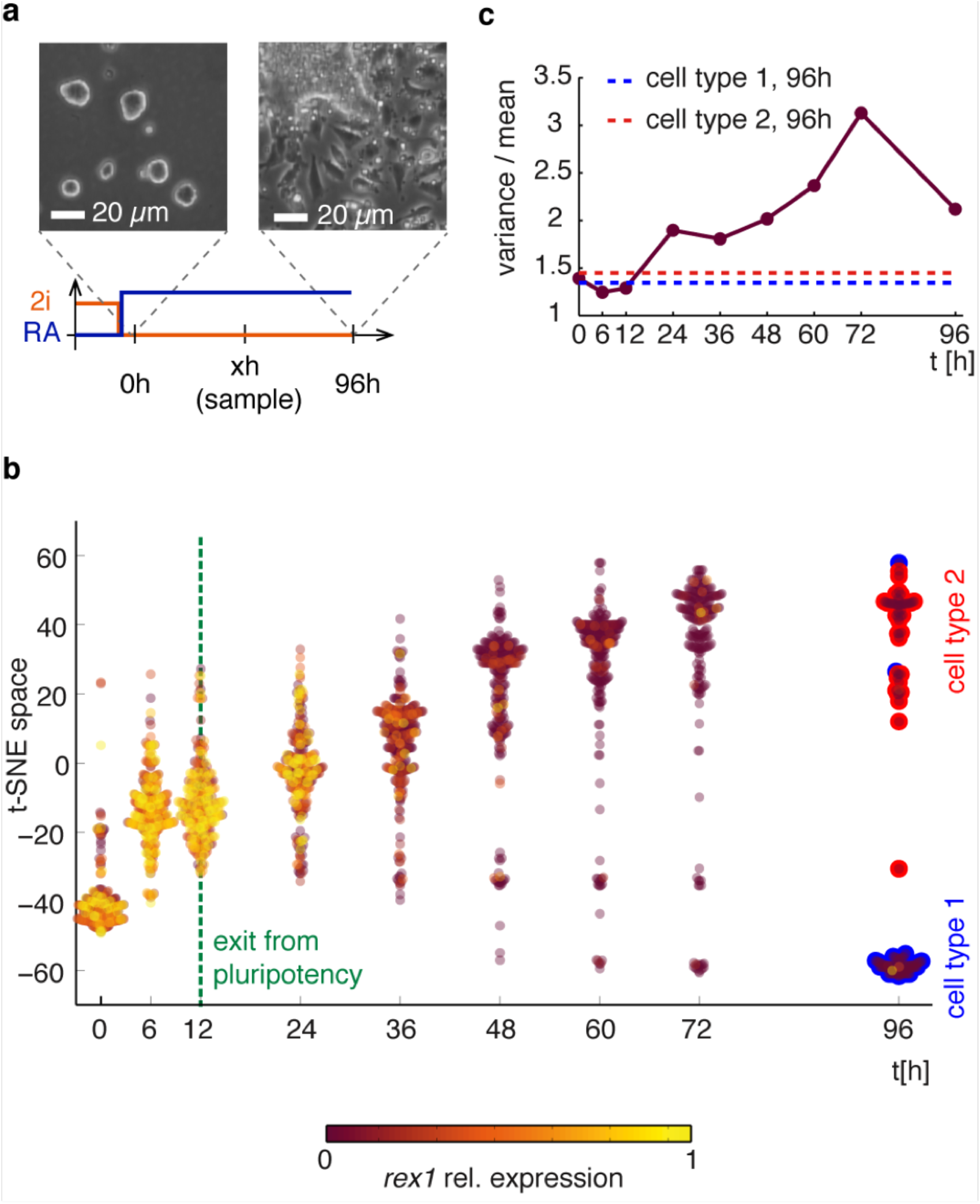
Single-cell RNA-seq revealed an RA driven bifurcation of mESCs after the exit from pluripotency. **a**, Scheme of the differentiation protocol with phase contrast images of cells growing in 2i medium (0 h) and after 96 h of exposure to 0.25 μM RA in N2B27 medium. **b**, t-SNE mapping of single-cell expression profiles. The single-cell RNA-seq data (SCRB-seq) for all cells and time points were mapped on a one-dimensional t-SNE space, which preserved local similarity between expression profiles, while reducing dimensionality. Each data point corresponds to a single cell. Data points for individual time points are shown in violin plots to reflect relative frequency along the t-SNE axis. The color of each data point indicates *rex1* expression (relative to maximum expression across all cells). For the 96 h time point, two robust clusters (found by k-means clustering and stability analysis) are indicated with red or blue edges, respectively. **c**, Single-cell gene expression variability quantified as the variance relative to the mean (Fano factor). The Fano factor of individual genes was averaged over all significantly variable genes. Expression was measured with SCRB-seq. Dashed lines indicate the average Fano factor calculated using only cells in one of the two clusters, or cell types after 96 h of RA exposure (see **b**).

**Figure 2.**
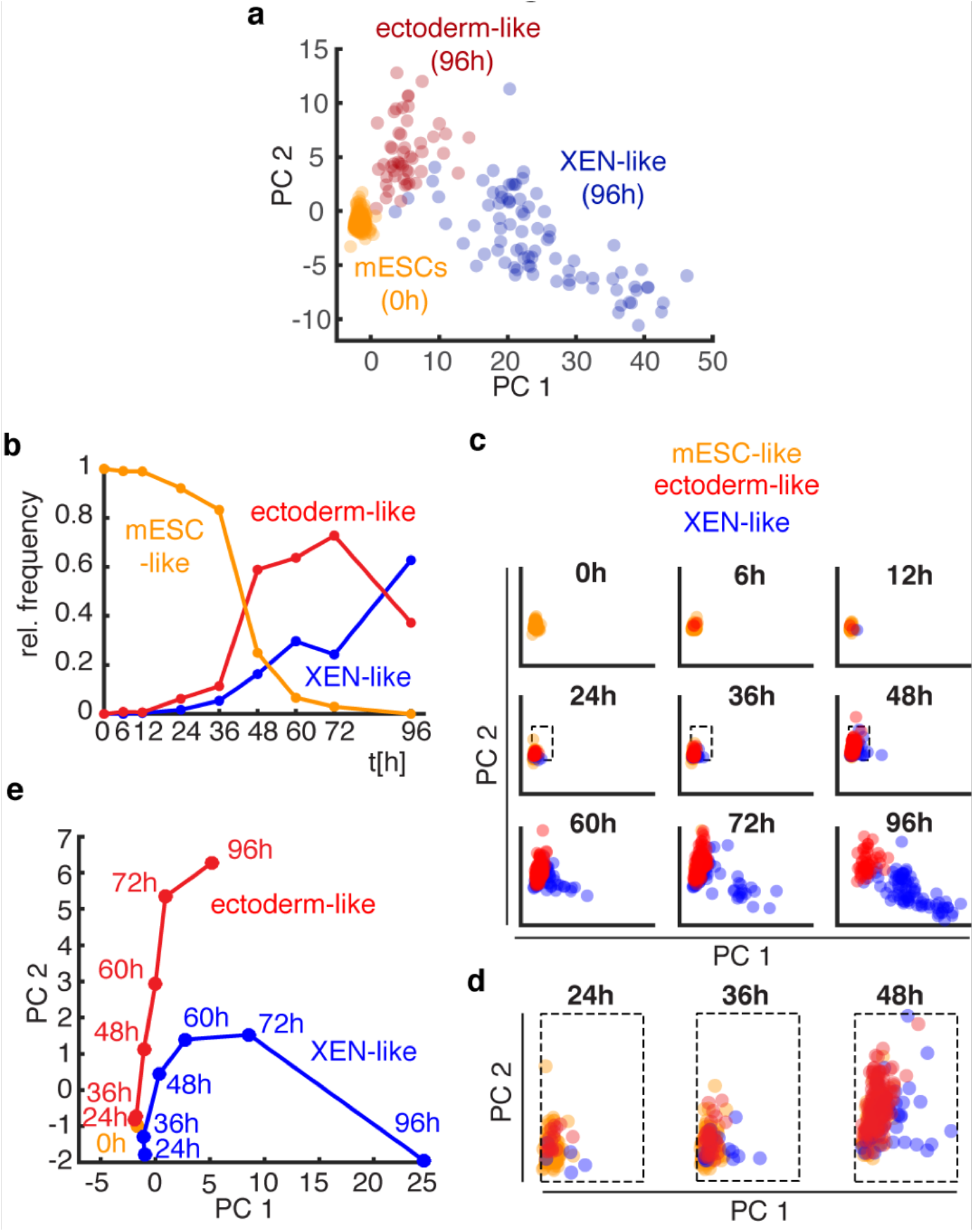
mESCs showed gradual adoption and divergence of lineage specific expression profiles after the exit from pluripotency. **a**, Principal component analysis of single-cell expression profiles of mESCs and after 96 h of RA exposure. Principal components were calculated across all cells and time points. Cells were placed in the space of the first two principal components (PC 1 and PC 2). Each data point corresponds to a single cell. Two robust clusters identified by k-means clustering and stability analysis are shown in red (ectoderm) and blue (XEN), respectively. mESCs are shown in yellow. **b**, Relative frequencies of cells classified as mESC-like, ectoderm- or XEN-like. Classification was based on Pearson correlation between expression profiles of individual cells and mean expression profiles of mESCs at 0 h or ectoderm-like and XEN-like cells after 96 h of RA exposure. An individual cell is identified with the cell type with which it is most strongly correlated. **c**, Principal component analysis of single-cell expression profiles. Principal components were calculated across all cells and time points. Cells measured at the indicated periods of RA exposure were placed in the space of the first two principal components. Each data point corresponds to a single cell. Cells were classified as mESC-like (orange), ectoderm-like (red) and XEN-like (blue) as described in the legend of panel **b**. The areas enclosed by dashed rectangles are shown in **d**. **d**, Same data as in **c** for three select time points (24h, 36h and 48h), zoomed in on the areas indicated by dashed rectangles in **c. e**, Average movement of ectoderm- and XEN-like cells in the principal component space during RA differentiation. The positions of cells of the same type were averaged at the indicated time points.

To quantify the divergence of expression profiles and identify the cell types represented by the two clusters, we used principal component analysis (PCA, Fig. 2a). We found that the first principal component (PC 1) was primarily composed of established markers for the XEN lineage (*sparc*, *col4a1*, *lama1*, *dab2*), while PC 2 comprised markers of ectodermal (neural) development (*prtg*, *mdk*, *fabp5*, *cd24*) (Supplementary Fig. 3c-d). Accordingly, we identified cell type 1 at 96 h as XEN-like and cell type 2 as ectoderm-like (Fig. 1b). Next, we sought to apply these classifications to the transitory time points post-pluripotency and pre-commitment. We classified cells at all time points as mESC-like, ectoderm-like and XEN-like (Fig. 2 b), which revealed a bifurcation into XEN-like or ectoderm-like expression profiles after around 24 h of RA exposure. This matched the phenomenological bifurcation in the t-SNE mapping (Fig. 1b). However, the majority of cells (roughly 60%) switched from an mESC–like, but no longer pluripotent state to a XEN-or ectoderm-like transcriptional program between 36 h and 48 h (Fig. 2 b). We concluded that cells first adopted lineage specific transcriptional programs 24 h to 36 h after the exit from pluripotency at 12 h. This suggested that some cells were not stably committed during this period. Interestingly, we found biases in the differentiation outcome depending on the timing of exit from pluripotency. Cells that down-regulated the mESC expression profile early (before 36 h) were biased towards the ectoderm lineage, while cells that exited the pluripotent state late (after 48h) adopted a XEN-like transcriptional program. This observation indicated that commitment to the two lineages did not occur simultaneously and that the lineage decision was initially biased towards ectoderm. We were further wondering whether the expression profiles of early XEN- and ectoderm-like cells were initially similar and, if so, when they diverged. We found that lineage specific expression profiles were established in a gradual, non-linear fashion: average XEN- and ectoderm expression states moved in a similar direction until 60 h after which they diverged more quickly (Fig. 2c-e). The similarity of the two lineages at the beginning of the bifurcation suggested that the cells were not fully committed to either lineage during this phase. Consequently, they might still be susceptible to a change in lineage cues during this phase.

**Figure 3.**
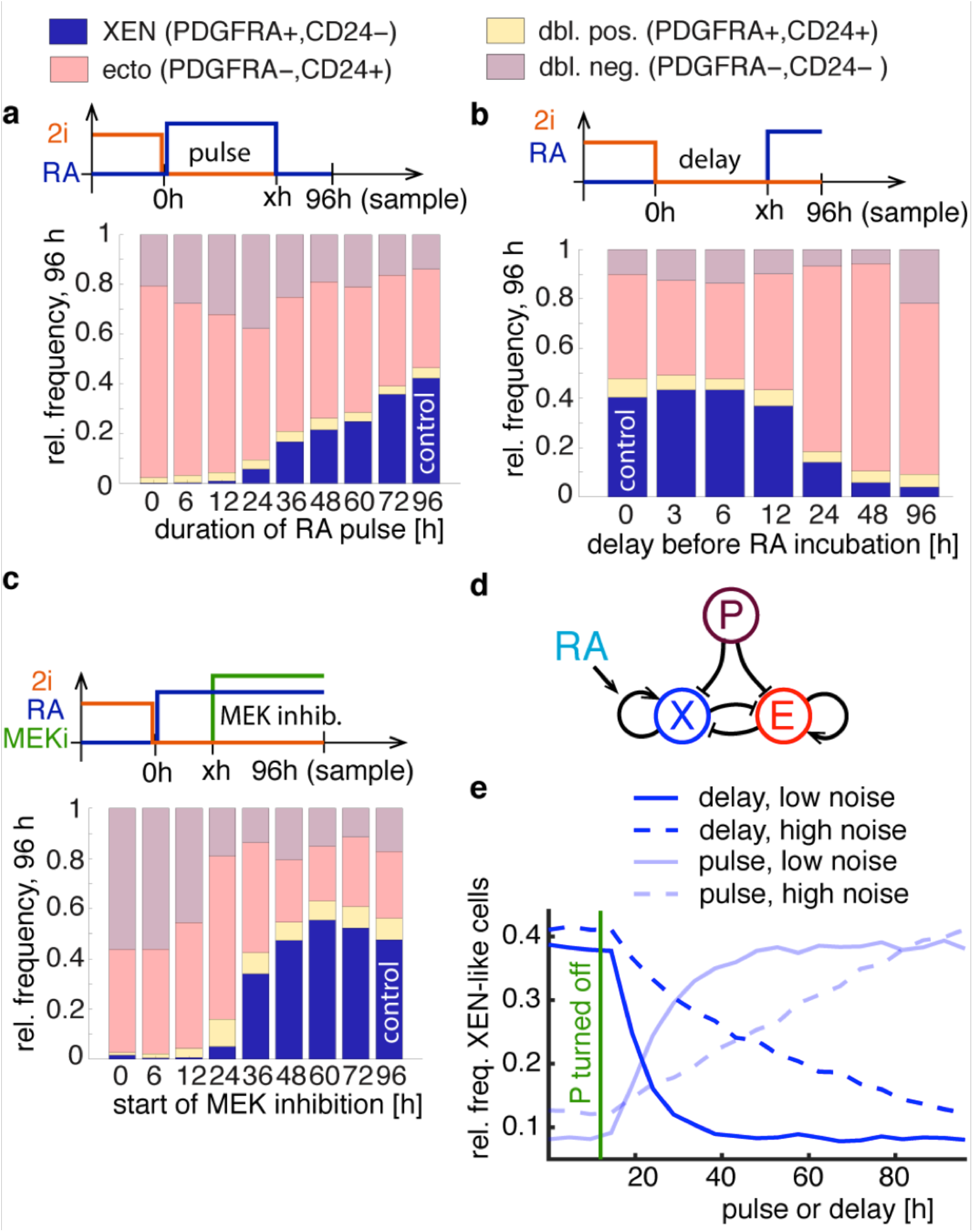
Susceptibility to signaling inputs was highly dynamic around the exit from pluripotency. **a,b,c** Fractions of cells classified as XEN-like, ectoderm-like, double positive and double negative after 96 h, based on *cd24* and *pdgfra* expression. Expression of the two markers was measured by antibody staining and flow cytometry. **a**, Cells were pulsed with 0.25 μM RA for x h and subsequently differentiated in basal medium (N2B27) complemented with an RA receptor antagonist. **b**, Cells were first incubated with basal medium (N2B27) for x h and then exposed to 0.25 μM RA for the remainder of the time course. **c,** Cells were incubated with 0.25 μM RA for x h after which 0.5 μM PD0325901 (MEK inhibitor) was added for the remainder of the time course. **d**, Schematic representation of a minimal gene regulatory network that can produce a lineage bifurcation ^20^. Pointy arrows indicate (auto-)activation;; blunted arrows indicate repression. E and X represent expression of ectoderm-like and XEN-like transcriptional programs, respectively. P stands for the pluripotency network. RA increases the auto-activation of the XEN program. **e**, Results of the stochastic simulations of the network shown in **d**. The relative frequency of XEN-like cells after 96 h is shown versus the length of an RA pulse or the length of the delay before RA exposure is started. In all cases the pluripotency network is turned off after 12 h. Simulations were run with low gene expression noise (solid lines) or high gene expression noise (dashed lines). See Supplementary Fig. 6b for exemplary trajectories.

To assay susceptibility directly, we next modulated the time of RA exposure during the differentiation process. We applied a precisely defined pulse of RA by first exposing the cells to RA for a defined period of time and then switching to a highly potent pan-RA receptor antagonist ^15^ (Fig. 3a). Cell type frequencies were quantified after 96 h using antibody staining of established surface markers for ectoderm (*cd24* ^16^) and XEN (*pdgfra* ^17^) (Supplementary Fig. 3a-b). These experiments showed that the RA pulse had to be applied for more than 12 h for XEN-like cells to appear. Longer pulses resulted in a gradual increase of the XEN-like fraction. A 36 h long pulse of RA resulted in 20% XEN-like cells at the 96 h time point, roughly half of what we found after uninterrupted RA exposure (Fig. 3a). This indicated that even after 36 h of RA exposure and significant downregulation of the pluripotency network many cells were not yet stably committed and XEN specification continued to depend on RA-signaling. We also wanted to establish when cells lost their ability to respond to RA signaling. To this end we first differentiated the cells in basal (N2B27) medium and started RA exposure after a defined time period. When RA exposure was delayed by up to 12 h, we did not observe any difference in the lineage distribution at the 96 h time point (Fig. 3b). Thereafter, for longer periods of RA-delay, we found that the fraction of XEN-like cells declined. This observation demonstrated that the cells gradually lost their susceptibility to RA after the exit from pluripotency. Taken together, the pulsed and delayed RA exposure experiments revealed a transient phase of about 24 h – 36 h after the exit from pluripotency, during which cells were maximally susceptible to external signaling cues to inform their lineage decision.

To generalize these results beyond RA exposure we next focused on signaling pathways that play pivotal roles in pluripotency and differentiation: we differentiated mESCs with RA in the presence of a MEK inhibitor (MEKi, PD0325901), which abrogates MAPK/Erk signaling; a GSK3 inhibitor, which effectively stimulates Wnt signaling (GSK3i, CHIR99021) or LIF, which activates the JAK/Stat pathway (Supplementary Fig. 4c). These signaling molecules are components of the defined 2i media and are known to prevent differentiation while stabilizing the pluripotent state. The presence of GSK3i or LIF led to an overall reduction of differentiated cells, consistent with their role in stabilizing pluripotency. Addition of MEKi alone, however, led to a specific reduction of the XEN-like subpopulation, in agreement with previous results ^18 19^. This effect was unlikely due to interference with RA signaling since increasing RA concentration did not reverse the effect (Supplementary Fig. 4d). Timed abrogation of MAPK/Erk signaling by MEKi mirrored the RA pulse experiments in terms of the effect on the lineage decision: at least 24 h of uninterrupted MAPK/Erk signaling was necessary for XEN-like cells to occur. Longer durations of MAPK/Erk signaling resulted in an increase in the XEN-like subpopulation (Fig. 3c). This effect plateaued after 48 h, which suggested that XEN-like cells then became independent of MAPK/Erk signaling and thus stably committed.

**Figure 4.**
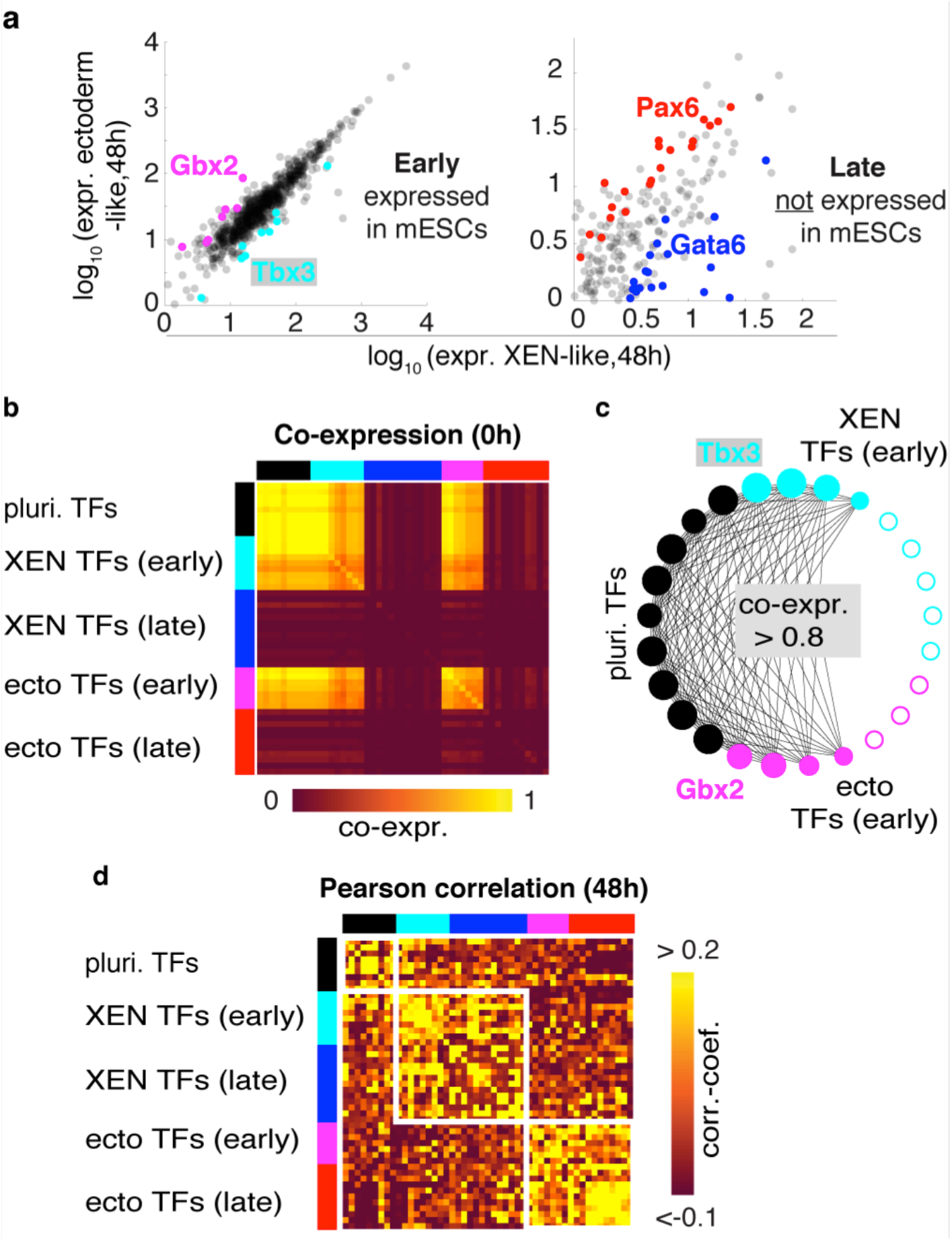
Distinct co-expression and correlation patterns identified two classes of lineage specific transcriptional regulators. **a**, Expression of transcriptional regulators in ectoderm-like and XEN-like cells identified in the SMART-seq data set. Genes that were significantly differentially expressed after 48 h of RA exposure are shown in red or pink (overexpressed in ectoderm-like cells) and blue or cyan (overexpressed in XEN-like cells), respectively. The two panels contain genes, which are present in the pluripotent state (early, left panel) or absent in the pluripotent state (late, right panel) A list of all identified factors is given in Supplementary Fig. 8b. **b**, Network of transcription factor co-expression in the pluripotent state. The gene set comprised the differentially expressed transcriptional regulators identified here (see **a**), as well as pluripotency related transcription factors listed in ^30^ (see Supplementary Fig. 8b). Co-expression was calculated using gene expression measured by SMART-seq. Co-expression of two genes was quantified as the fraction of cells in which the expression of both genes exceeded a certain threshold value (see Methods). **c**, Co-expression network in the pluripotent state. Two factors are connected by an edge if their co-expression exceeds 0.8. The gene set comprised XEN specific factors (cyan nodes) and ectoderm specific factors (pink nodes) that are expressed in the pluripotent state (early factors), as well as pluripotency factors listed in ^30^ (black nodes). The radius of solid nodes is proportional to the number of connections to other nodes. Nodes without any connections are depicted as open nodes. **d**, Pearson correlation between a set of core transcriptional regulators after 48 h of RA exposure. The gene set is the same as in **b**. Pearson correlation was calculated using gene expression measured by SMART-seq. Pearson correlation coefficients exceeding the range [−0.1,0.2] are shown in the same color as the respective extreme values of that range.

In summary, the signaling experiments showed that cells acquired susceptibility to lineage specifying cues immediately after the exit from pluripotency and remained susceptible to a change in signaling input for about 24 h - 36 h (see schematic in Supplementary Fig. 4e).

The sudden increase in gene expression variability at the beginning of the susceptible phase (Fig. 1c) led us to hypothesize that gene expression noise might have an important impact on the dynamics of commitment ^5 4^. Since the strength of such noise is not amenable to experimental manipulation we instead sought to develop a simple phenomenological model that would explain the existence of a transient phase of susceptibility. We were also wondering what might determine the length of this phase. Our model is based on a minimal gene regulatory network (GRN) that has been used before to describe lineage bifurcations ^19,20^ (Fig. 3d, Supplementary Fig. 5a). Briefly, the GRN is comprised of two lineage-specific, auto-activating expression programs that mutually repress each other. This GRN can produce two stable attractors, corresponding to the two cell lineages. Here, we added repression of both lineages by the pluripotency network to model the pluripotent state. Consistent with our data, we assumed that the pluripotency program is turned off after 12 h. Since RA exposure is necessary for the occurrence of XEN-like cells (Fig. 3b), we modeled the effect of RA as increased auto-activation of the XEN program. Stochastic simulations of this 3-state GRN reproduced a transient phase of susceptibility to RA exposure (Fig. 3e). In this phase, both lineage specific programs are co-expressed in individual cells and slight (stochastic) imbalances between the two programs can bias cells to one or the other lineage (Supplementary Fig. 5b). In a subsequent phase, gene expression profiles diverge and approach the stable attractor states. Consequently, cells lose susceptibility to a change in RA signaling input. Importantly, increased gene expression noise led to a longer phase of susceptibility: in the presence of high noise, cells that are already close to one of the attractors can still escape to the other attractor, when the signaling input is switched.

**Figure 5.**
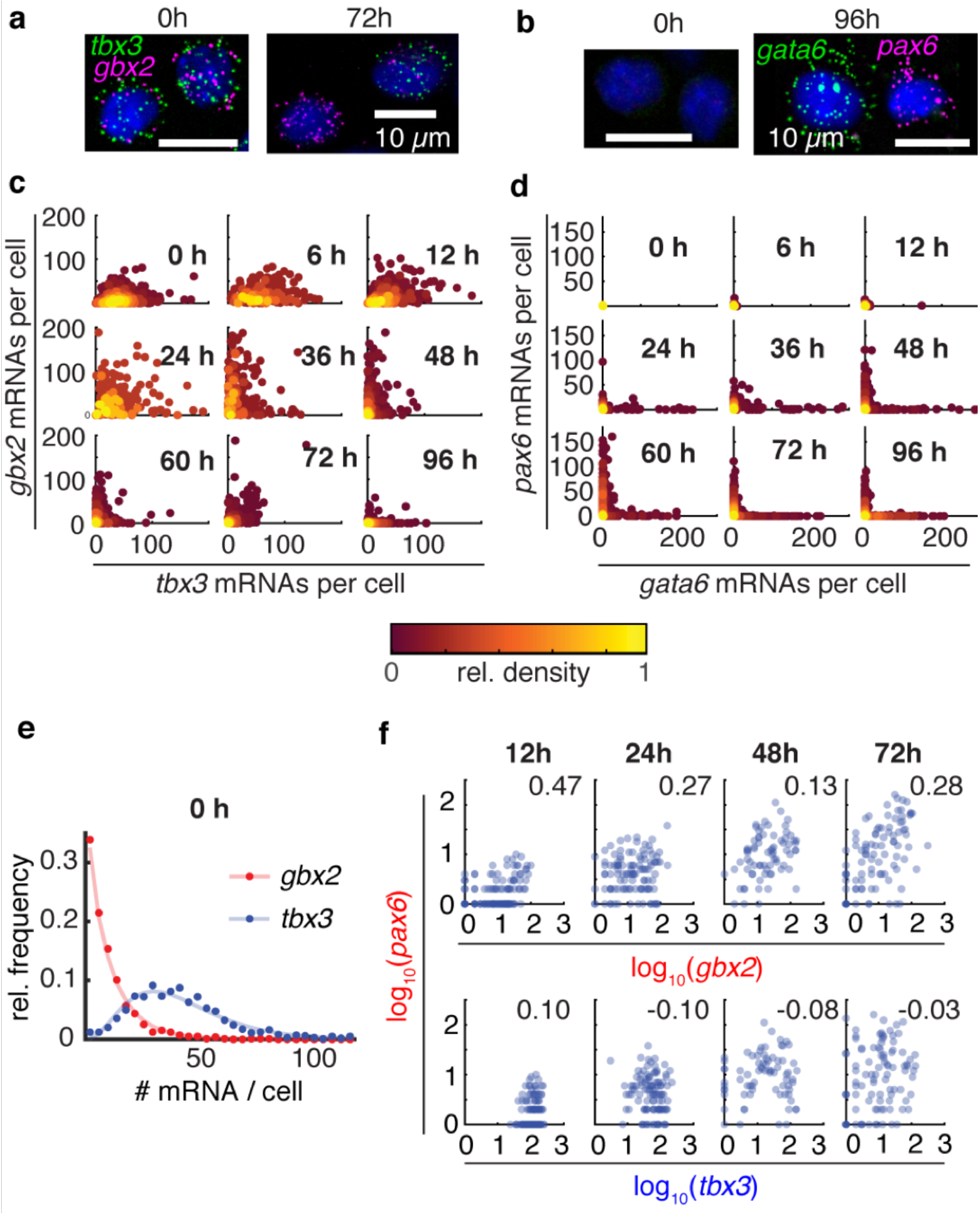
smFISH confirmed distinct expression patterns of exemplary transcription factors. **a**, Fluorescence images of smFISH for *gbx2* and *tbx3* in mESCs (0 h) and after 72 h RA exposure. Each diffraction limited dot corresponds to a single mRNA molecule. Hoechst staining of nuclei is shown in blue. **b**, Fluorescence images of smFISH for *pax6* and *gata6* in mESCs (0 h) and after 96 h RA exposure. Each diffraction limited dot corresponds to a single mRNA molecule. Hoechst staining of nuclei is shown in blue. **c**, Scatter plots of the number of *gbx2*and *tbx3* mRNAs per cell measured by smFISH. Each data point is a single cell. Color indicates the local density of data points. The number of shown cells measured at a certain time point ranges between 224 and 983. **d**, Scatter plots of the number of *pax6* and *gata6* mRNAs per cell measured by smFISH. Each data point is a single cell. Color indicates the local density of data points. The number of shown cells measured at a certain time point ranges between 293 and 570. **e**, Distribution of the *tbx3* and *gbx2* transcripts in individual mESCs as measured by smFISH. Both data sets are fit by a Gamma distribution (*tbx3,* R^2^ = 0.94, solid blue line;; *gbx2*, R^2^ = 0.99, solid red line). **f**, Scatter plots of the number of mRNAs per cell for *gbx2* and *tbx3* vs*pax6* measured by smFISH. Each data point is a single cell. Cells were exposed to RA for 12 h, 24 h, 48 h and 72 h, respectively, as indicated above each column of panels. The number in each panel is the Pearson correlation between the genes plotted in the respective panel.

Based on our experimental observations as well the phenomenological model, we now suggest a two-phase differentiation process. In the first phase, the lineage-biasing phase, right after exit from pluripotency, XEN and ectoderm programs are co-expressed in individual cells. In this phase, imbalances between the two programs can bias the lineage decision and cells are susceptible to lineage cues. Subsequently, the cells enter the second phase, the lineage-committing phase. In this phase the two lineage specific programs become mutually exclusive and cell states become independent of the original lineage specifying cue. We next set out to validate our model by finding transcription factors that might constitute a mutually-repressive GRN. Consequently, we wanted to focused on transcription factors that show lineage specific expression at the beginning of the lineage-committing phase, around 48 h. Since transcription factors are typically lowly expressed, they are not well-represented in the SCRB-seq data set. We collected another single-cell RNA-seq data set using SMART-seq2 ^21^ at four early RA differentiation time points (0 h, 12 h, 24 h and 48 h). After identification of XEN-like or ectoderm-like cells at the 48 h time point (Supplementary Fig. 6a) we found 55 transcription factors to be differentially expressed between the two lineages, at that time point (Fig. 4a, Supplementary Fig. 6b). 16 of those transcription factors (dubbed “early” transcription factors) were present throughout the differentiation time course. Consistent with our model, early factors were broadly co-expressed in individual cells at the beginning of the time course (Fig. 4b and Supplementary Fig. 6c). Compared to canonical pluripotency factors in the pluripotent state, early factors showed a smaller level of co-expression with each other, in particular if they belonged to different lineages (Fig. 4b, c). Individual cells thus expressed varying ratios of XEN and ectoderm specific early factors during the lineage-biasing phase, as observed in our simulations. Over time, co-expression of XEN and ectoderm specific early factors declined but they never became completely mutually exclusive (Supplementary Fig. 6c). We speculated that other transcription factors might be up-regulated in lineage biased cells and take over lineage specification from the early factors. Indeed, 39 of the identified differentially expressed transcription factors (dubbed “late” transcription factors) were not significantly expressed at the beginning of the time course (Fig. 4b). These late factors were overall positively correlated with early factors of the same lineage and anti-correlated with factors of the opposing lineage (Fig. 4d and Supplementary Fig. 6c). This correlation pattern suggested that early factors have a role in lineage biasing, whereas late factors are involved in lineage commitment.

To further validate the notion of a handover between early and late factors, we next focused on four transcription factors, chosen based on their reported function for the specification of ectoderm (*gbx2* ^22^ (early), *pax6* ^23^ (late)) and extraembryonic endoderm (*tbx3* ^24^ (early), *gata6* ^25^ (late)). In agreement with their reported roles we found these factors to be differentially expressed in ectoderm-like and XEN-like cells, respectively, in our SCRB-seq data set (Supplementary Fig. 6d). To quantify correlation patterns with high precision we used single-molecule FISH (smFISH ^26^) due to its superior performance compared to single-cell RNA-seq (Supplementary Fig. 7a). We measured the expression of the early factors (Figs. 5 a,c) or the late factors (Figs. 5b,d) simultaneously and quantified co-expression at all time points (Supplementary Fig. 7b-d). In agreement with the SMART-seq data, co-expression of early factors was highest in the pluripotent state and declined after exit from pluripotency (Fig. 5c, Supplementary Fig. 7c). Importantly, mESCs expressed the early factors at highly variable ratios: 30% of mESCs did not express the early ectoderm factor *gbx2* at a significant level, while almost all cells expressed the early XEN marker *tbx3* (Fig.5e). smFISH further confirmed that late factors were only sporadically expressed before the exit from pluripotency but strongly up-regulated in separate subpopulations thereafter. These subpopulations likely corresponded to lineage-committed cell states (Fig. 5d and Supplementary Fig. 7d-e). A simultaneous measurement of the early ectoderm factor *gbx2* and and the late ectoderm factor *pax6* also provided direct support of the suggested handover between transcription factor classes: the two factors were positively correlated throughout the time course, even before the exit from pluripotency (Fig. 5f). We speculate that the cells with the highest expression of the early factor were biased towards the ectoderm lineage, and consequently up-regulated the late ectoderm factor. All in all, the smFISH experiments clearly confirmed differences in the expression dynamics of early and late factors, supporting their classification and suggested functions.

## DISCUSSION

In summary, we have leveraged a recently developed high-throughput single-cell transcriptomics method to dissect the dynamics of lineage bifurcation and commitment in RA driven differentiation of mESCs with high temporal resolution. A recently published study by Klein et al. used single-cell RNA-seq to characterize mESC differentiation by LIF withdrawal ^27^. Klein et al. also found a XEN-like subpopulation but due to the low temporal resolution of their experiment (samples at 2d, 4d and 7d post LIF-withdrawal) detailed bifurcation dynamics were not revealed. Concerning the susceptibility to lineage cues our results have interesting parallels with an earlier report by Turner et al. ^28^. That study analyzed a lineage decision between the neuroectoderm and primitive streak lineage in mESCs. Turner et al. showed that the susceptibility to primitive streak inducing cues was strongly dynamic, albeit on a time scale of several days. The authors interpreted their results in terms of a primary neuroectodermal fate and a delayed acquisition of potency to adopt endomesodermal fates. Similarly, here we observed transient susceptibility to RA signaling and the appearance of ectoderm-like cells before XEN-like cells. However, we explain the gradual occurrence of committed cells by the presence of gene expression noise. This model was inspired by the observation of a sharp increase in gene expression variability at the exit from pluripotency. This increase might result from a loss of tight gene regulation, which is necessary for acquiring susceptibility to lineage cues. The impact of noise in the context of lineage bifurcations was recently addressed in a publication by Marco et al. ^29^. In that study the authors focused on the ability of noise to destabilize committed cell states. Here we extended their considerations to the effects of noise on commitment dynamics by stochastic simulations of a minimal GRN. A similar GRN had been used successfully before in a report by Schröter et al., studying induction of the XEN lineage by exogenous *gata4* expression ^19^. The influence of noise can be understood intuitively by examining our simulated gene expression trajectories (Supplementary Fig. 5b). In the case of high gene expression noise, trajectories could switch more easily between the basins of attraction of the two attractors. Consequently, cells only became truly committed when they were very close to one of the attractors, towards the end of the simulation period. Hence, tuning the strength of gene expression noise could be a mechanism by which cells influence the length of the susceptible phase. Conversely, *in vitro* differentiation protocols might benefit from eliminating sources of extrinsic noise (like fluctuating culture environments, for example).

Our study also identified early-expressed lineage specific transcription factors that are heterogeneously expressed in the pluripotent state and thus have a potential role in biasing the lineage decision. Importantly, the two factors we studied in detail, *gbx2* and *tbx3*, were previously determined to be part of an essential pluripotency network ^30^. It has been suggested before that some pluripotency genes are also involved in lineage specification ^31 32^. Thomson et al. showed that *sox2* and *oct4* promote the neuroectodermal and mesendodermal lineage, respectively ^31^. Future research will have to show whether *gbx2* and *tbx3* have similar roles for the ectoderm and XEN lineage, respectively. In fact, for *tbx3* there is some evidence for a dual function in self-renewal and XEN specification ^24^. The observed correlation between *gbx2* and *pax6* suggests a function of *gbx2* in ectoderm specification. The long-tail distribution of *gbx2* hints at infrequent transcriptional bursting and possibly distinct subpopulations ^33^. The causal relationship between gbx2 and *pax6* and the functional relevance of the *gbx2* high subpopulations will be explored in a future study. Late-expressed lineage specific transcription factors, like *pax6* and *gata6*, which were not expressed in the pluripotent state, have a potential role in lineage commitment. They can thus serve as *bona fide* lineage markers.

Transient phases of susceptibility to lineage cues, such as the one characterized in this study, might be valuable windows of opportunity for the control of lineage decisions. We speculate that exit from a pluripotent cell state is necessarily followed by a phase of instability, likely generalizing our findings to most differentiation systems. Based on our results we would like to propose tentative transcriptional signatures of such phases: 1. down-regulation of pluripotency factors (Supplementary Fig. 1b), 2. a sudden increase in single-cell gene expression variability (Fig. 1c), 3. slowly diverging lineage specific expression patterns (Fig. 2e), 4. co-expression of early-expressed (lineage-biasing) transcription factors (Fig. 5c, Supplementary Fig. 7c) and 5. sporadic expression of late-expressed (lineage-committing) transcription factors (Fig. 5d, Supplementary Fig. 7d). We hope that these results will be a stepping stone towards finding more efficient ways to guide lineage decisions.

## MATERIALS & METHODS

### Cell culture

E14 or V6.5 mouse embryonic stem cells were grown in modified 2i medium ^9^: DMEM/F12 (Life technologies) supplemented with 0.5× N2 supplement, 0.5× B27 supplement, 4mM L-glutamine (Gibco), 20 μg/ml human insulin (Sigma-Aldrich), 1× 100U/ml penicillin/streptomycin (Gibco), 1× MEM Non-Essential Amino Acids (Gibco), 7 μl 2-Mercaptoethanol (Sigma-Aldrich), 1 μM MEK inhibitor (PD0325901,Stemgent), 3 μM GSK3 inhibitor (CHIR99021, Stemgent), 1000 U/ml mouse LIF (ESGRO). Cells were passaged every other day with Accutase (Life technologies) and replated on gelatin coated tissue culture plates (Cellstar, Greiner bio-one).

### Differentiation

Prior to differentiation cells were grown in 2i medium for at least 2 passages. Cells were seeded at 2.5 × 10^5^ per 10 cm dish and grown over night (12 h). Cells were then washed twice with PBS and basal N2B27 medium (2i medium without the inhibitors, LIF and the additional insulin) supplemented with all-trans retinoic acid (RA, Sigma-Aldrich). RA concentration was 0.25 μM unless stated otherwise. Spent medium was exchanged with fresh medium after 48 h.

For the RA pulse experiments (Fig. 3a) cells were first differentiated with 0.25 μM RA for the indicated amounts of time, washed three times with PBS and cultured in basal medium with 2.5 μM of the RA receptor antagonist AGN 193109 (sc-210768, Santa Cruz Biotechnology). At this concentration this antagonist completely inhibits signaling through all-trans retinoic acid ^15^.

For the differentiation under perturbation of various signaling pathways (Supplementary Fig. 4c) we used the MEK inhibitor PD0325901 (Stemgent, standard concentration 1 μM or dilutions thereof), GSK3 inhibitor CHIR99021 (Stemgent, standard concentration 3 μM or dilutions thereof) or mouse LIF (ESGRO, 1000 U/ml). For the experiments with MEK inhibition shown in Fig. 3c and Supplementary Fig. 4d we used PD0325901 at a concentration of 0.5 μM.

### Long-term culture of differentiated cells

After sorting differentiated cells were replated on poly-D-lysine and laminin coated tissue culture dishes in basal (N2B27) medium complemented with 20 ng/ml mouse EGF (E5160, Sigma-Aldrich) and 10 ng/ml mouse FGF2 (SRP4038-50UG, Sigma-Aldrich). Ectoderm-like cells were propagated by dissociation with accutase (Life Technoliges) and replating under identical conditions every 3-4 days. Floating aggregates of XEN-like cells were propagated in suspension in uncoated plastic petri dishes. Aggregates were not dissociated but the medium was refreshed roughly every 4 days.

### Antibody staining

We used the following antibodies: APC Rat Anti-Mouse *cd24* (BD Bioscience, 562349), PE Rat Anti-Mouse *cd24* (BD Bioscience, 553262), Anti-Mouse CD140a (PDGF Receptor a) FITC (eBioscience,17-1407), Anti-Mouse CD140a (PDGF Receptor a) APC (eBioscience,17-1401), all at a dilution of 1:1000. Cells growing in 6-well plates were washed once with PBS and then incubated in a volume of 500 μl of basal (N2B27) medium with antibodies for 30 min at 37 ^°^C, in the dark. Subsequently, cells were washed once with PBS, 300 μl Accutase (Life Technologies) was added and cells were gently dissociated by pipetting up and down. After adding 600 μl of basal medium the cell suspension was loaded on a flow cytometer (LSR II, BD Bioscience). Cells growing in 10 cm dishes were first dissociated and incubated in 1 ml medium with the same incubation conditions and antibody concentrations as for adherent cells. After staining in solution, cells were spun down, the supernatant was removed and cells were resuspended in 1 ml of basal medium before flow cytometry.

### Colony formation assay

Cells were differentiated with or without RA as described above for various amounts of time and then replated at a density of 5 × 10^4^ cells/well in a gelatinized 6-well tissue culture plate in 2i medium. Colonies were grown for 2 additional days, washed twice with PBS and then imaged in PBS. Remaining colonies were counted automatically by a custom made image analysis script written in MATLAB. The number of surviving colonies was normalized to the first data point (replating of untreated cells growing in 2i).

### Measurement of cell cycle phases

Cells growing on gelatinized tissue culture dishes were washed twice with PBS, detached with Accutase (Life technologies) and resuspended in full medium. Formaldehyde was added to the cell suspension to a final concentration of 4%. Cells were incubated for 12 min at room temperature while being rotated and then spun down for 3 min at 1,000 rpm. Subsequently cells were permeabilized at least over night in 70% ethanol. Cells were stained with Hoechst 33342 in PBS for 1 h and fluorescence measured on a flow cytometer (LSR II, BD Biosciences). The Dean-Jet-Fox model ^34^ was fit to histograms of the fluorescence signal to determine the relative lengths of the cell cycle phases reported in Supplementary Fig. 1e.

### Single cell isolation for SCRB-seq

For each differentiation time point cells were harvested and medium removed by spinning for 5 min at 1000 rpm. RNA was stabilized by immediately resuspending the pelleted cells in RNAprotect Cell Reagent (Qiagen) and RNaseOUT Recombinant Ribonuclease Inhibitor (Life Technologies) at a 1:1000 dilution. Just prior to fluorescence-actived cell sorting (FACS), the cells were diluted in PBS and stained for viability using Hoechst 33342 (Life Technologies). 384-well SBS capture plates were filled with 5μl of a 1:500 dilution of Phusion HF buffer (New England Biolabs) in water and individual cells were then sorted into each well using a FACSAria II flow cytometer (BD Biosciences) based on Hoechst DNA staining. After sorting, the plates were immediately sealed, spun down, cooled on dry ice and then stored at −80^°^C.

### SCRB-Seq of sorted single cells

Frozen cells were thawed for 5 minutes at room temperature and cell lysis was enhanced by a treatment with proteinase K (200 μg/mL;Ambion) followed by RNA desiccation to inactivate the proteinase K and simultaneously reduce the reaction volume (50 ^°^C for 15 min in sealed plate, then 95 ^°^C for 10 min with seal removed).

To start, diluted ERCC RNA Spike-In Mix (1 μl of 1:10^7^; Life Technologies) was added to each well and the template switching reverse transcription reaction was carried out using Maxima H Minus Reverse Transcriptase (Thermo Scientific), our universal adapter E5V6NEXT (1 pmol, Eurogentec):

5’-iCiGiCACACTCTTTCCCTACACGACGCrGrGrG-3’

where iC: iso-dC, iG: iso-dG, rG: RNA G, and our barcoded adapter E3V6NEXT (1 pmol, Integrated DNA Technologies):

5′-/5Biosg/ACACTCTTTCCCTACACGACGCTCTTCCGATCT[BC6]N10T30VN-3′

where 5Biosg = 5’ biotin, [BC6] = 6bp barcode specific to each cell/well, N10 = Unique Molecular Identifiers. Following the template switching reaction, cDNA from 384 wells was pooled together, and then purified and concentrated using a single DNA Clean & Concentrator-5 column (Zymo Research). Pooled cDNAs were treated with Exonuclease I (New England Biolabs) and then amplified by single primer PCR using the Advantage 2 Polymerase Mix (Clontech) and our SINGV6 primer (10 pmol, Integrated DNA Technologies):

5’-/5Biosg/ACACTCTTTCCCTACACGACGC-3’

Full length cDNAs were purified with Agencourt AMPure XP magnetic beads (0.6x, Beckman Coulter) and quantified on the Qubit 2.0 Flurometer using the dsDNA HS Assay (Life Technologies). Full-length cDNA was then used as input to the Nextera XT library preparation kit (Illumina) according to the manufacturer’s protocol, with the exception that the i5 primer was replaced by our P5NEXTPT5 primer (5μM, Integrated DNA Technologies):

5’-

AATGATACGGCGACCACCGAGATCTACACTCTTTCCCTACACGACGCTCTTCCG*A*T*C*

T*-3’

where ^*^ = phosphorothioate bonds.

The resulting sequencing library was purified with Agencourt AMPure XP magnetic beads (0.6x, Beckman Coulter), size selected (300-800bp) on a E-Gel EX Gel, 2% (Life Technologies), purified using the QIAquick Gel Extraction Kit (Qiagen) and quantified on the Qubit 2.0 Flurometer using the dsDNA HS Assay (Life Technologies). Libraries were sequenced on Illumina Hiseq paired-end flow cells with 17 cycles on the first read to decode the well barcode and UMI, an 9 cycle index read to decode the i7 Nextera barcode and finally a 46 cycle second read to sequence the cDNA.

### RNA-seq on bulk samples

Populations of sorted cells were collected in RNAprotect, lysed in QIAzol (Qiagen) and total RNA was extracted and purified using Direct-zol RNA MiniPrep (Zymo Research). DGE libraries were prepared from 10 ng of extracted total RNA, using the protocol previously described for SCRB-seq with the exception of using more concentrated E3V6NEXT and E5V6NEXT (10 pmol).

### SCRB-seq and bulk RNA-seq read alignment

All second sequence reads were aligned to a reference database consisting of all mouse RefSeq mRNA sequences (obtained from the UCSC Genome Browser mm10 reference set: http://genome.ucsc.edu/), the mouse mm10 mitochondrial reference sequence and the ERCC RNA spike-in reference sequences using bwa version 0.7.4 4 with non-default parameter “-l 24”. Read pairs for which the second read aligned to a mouse RefSeq gene were kept for further analysis if 1) the initial six bases of the first read all had quality scores of at least 10 and corresponded exactly to a designed well-barcode and 2) the next ten bases of the first read (the UMI) all had quality scores of at least 30. Digital gene expression (DGE) profiles were then generated by counting, for each microplate well and RefSeq gene, the number of unique UMIs associated with that gene in that well. Python scripts implementing the alignment and DGE derivation are available from the authors upon request.

### SMART-seq sample preparation and read alignment

The single-cell SMART-seq2 libraries were prepared according to the SMART-seq2 protocol^21 35^ with some modifications ^36^. Briefly, total RNA from single cells sorted in lysis buffer was purified using RNA-SPRI beads. Poly(A)+ mRNA from each single cell was converted to _c_DNA which was then amplified. cDNA was subjected to transposon-based fragmentation that used dual-indexing to barcode each fragment of each converted transcript with a combination of barcodes specific to each single cell. Barcoded cDNA fragments were then pooled prior to sequencing. Sequencing was carried out as paired-end 2x25bp with 8 additional cycles for each index. Alignment of the reads and calculation of gene expression was done through the Tuxedo pipeline (Tophat, Cuffquand, Cuffnorm) ^37^. Gene expression was expressed as reads per kilobase exon model per million mapped reads (RPKM).

### Computational analysis of the bulk RNA-seq experiments

The bulk RNA-seq results were normalized by the total amount of reads per time point. Only those genes with non-zero mean were considered for further analysis. For k-means clustering of the temporal profiles we first determined the number of robust clusters. Stability analysis ^14^ indicated that there were 6 robust clusters (Supplementary Fig. 3a). We then performed gene ontology enrichment analysis using the DAVID bioinformatics resource ^38^ the results of which are summarized in Supplementary Fig. 3b. Only the clusters of monotonically upregulated genes showed significant enrichment for GO terms related to development, morphogenesis and differentiation. The heat maps of bulk RNA-seq data depict expression relative to *gapdh* expression (Supplementary Fig. 1a).

### Computational analysis of the SCRB-seq experiments

A histogram of the total number of UMIs detected per cell is shown in Supplementary Fig. 2a. To reduce the influence of technical noise we discarded cells with less than 2000 UMIs (red vertical line in Supplementary Fig. 2). This cutoff nearly minimized the upper bound of the counting error per gene (Supplementary Fig. 2b) estimated by

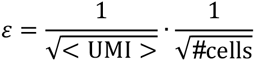

while not significantly reducing the number of detected genes (13720, Supplementary Fig. 2c) - defined as the number of genes, which had more than one UMI in more than one cell. Due to this cutoff 2451 out of 3456 measured cells were used for further analysis (Supplementary Fig. 2e). In individual cells with more than 2000 UMIs in total on average 850 genes were detected.

For all further analyses, except the calculation of Fano factors, the data was normalized in the following way to account for differences in efficiency of transcript recovery between wells: UMI counts were divided by the total number of UMI counts per cell and then multiplied by the median of total UMI counts across all cells growing in 2i medium. For the calculation of Fano factors (Fig. 1c) UMI counts were down-sampled to 2000 UMI counts per cell. This down-sampling procedure ensured that the contribution of counting error to the Fano factors was equal for all cells from all time points. To include only those genes, which exhibited significant, biological variability, we plotted the coefficient of variation (CV) of individual genes over all time points with respect to the mean expression level as well as the CVs of ERCC spike-ins with known abundance (Supplementary Fig. 2f). The increase in variability with decreased average expression reflected higher technical and counting noise for lowly expressed genes. We used the 829 genes, which had the 5% highest ratios of CV and the moving average of the CV for principal component analysis, k-means clustering and t-SNE mapping (see below).

To further characterize the performance of SCRB-seq we first compared SCRB-seq data averaged over cells for individual time points with bulk RNA-seq and found them to be strongly correlated (Supplementary Fig. 2g, Pearson correlation ρ = 0.75). We compared 100 randomly selected pairs of cells growing in 2i medium and found that SCRB-seq measurements of individual cells were strongly correlated (Supplementary Fig. 2h, Pearson correlation ρ = 0.63). By analysis of UMI counts of ERCC spike-in RNA we determined that UMI counts scaled approximately linearly with the spiked-in transcripts - the slope of a linear fit to the log-log plot of spike ins versus UMI counts was 0.78. The efficiency of transcript recovery as determined from the offset of that linear fit was about 0.9% (Supplementary Fig. 2i).

For principal component analysis (PCA) we considered genes, which belonged to the upregulated clusters (clusters 5 and 6, Supplementary Fig. 3a) and were among the most variable genes (Supplementary Fig. 2f). Prior to PCA expression profiles of individual genes were converted to z-scores using the average expression over all time points and the moving average of the coefficient of variation (Supplementary Fig. 2f) to preserve biological variability. PCA was performed with all cells across all time points and expression profiles of individual cells were then projected on the found principal components. The genes with the highest loadings in the first two principal components are listed in Supplementary Fig. 3c and their loadings are represented graphically in Supplementary Fig. 3d.

To discover clusters of cells we used k-means clustering including all 829 most variable genes and using (1 - Pearson correlation) as the distance metric. Cluster-wise assessment of stability ^14^ was used to determine the robustness of clusters. In particular, we calculated the Jaccard similarities between clusters found in bootstrapped samples. Clusterings resulting in Jaccard similarities close to 0.5 were considered unstable. In this way clusters were found for the 96 h time point. For earlier time point cells were classified according to similarity with the clusters found at 96 h or mESCs at 0h. In particular, we first calculated the mean expression profiles of mESCs, as well as the XEN-like and ectoderm-like subpopulations at 96 h. Then Pearson correlation was calculated between those average profiles and expression profiles of individual cells at earlier time points. A cell was classified as a particular cell type when the correlation with this particular cell type exceeded the correlation with all other cell types.

Gene expression of individual genes was represented in color by normalizing to the maximum expression per time point, linear histogram stretching (1^st^ to 99^th^ percentile) and subsequent linear mapping to a custom colormap (Supplementary Fig. 6d).

For t-distributed stochastic neighbor embedding (t-SNE) we considered genes, which were among the most variable genes (Supplementary Fig. 2f). Prior to t-SNE mapping profiles of individual genes were converted to z-scores using the average expression over all time points and the moving average of the coefficient of variation (Supplementary Fig. 2f) to preserve biological variability. One-dimensional t-SNE maps were computed using the MATLAB Toolbox for Dimensionality Reduction (v0.8.1 - March 2013) (^39^, L.J.P. van der Maaten, http://homepage.tudelft.nl/19j49/Matlab_Toolbox_for_Dimensionality_Reduction.html). Expression of *rex1* was represented in color by normalizing to the maximum expression, linear histogram stretching (0^th^ to 95^th^ percentile) and subsequent linear mapping to a custom colormap.

### Computational analysis of the SMART-seq experiments

Only cells with at least 200000 reads per cell were used. For all further analyses the data was normalized in the following way to account for differences in the total number of reads between samples: RPKM for individual genes were divided by the total number of RPKM per cell and then multiplied by the median of total RPKM across all cells growing in 2i medium. Cells with high expression of *cd24* or *pdgfra* were classified as shown in Supplementary Fig. 6a. Out of the 82 cells measured by SMART-seq at 48 h, 10 were considered XEN-like (*pdgfra* high) and 29 ectoderm-like (*cd24* high). To compute significance levels for gene expression in these subpopulations we used a null model that assumes that all cells were essentially identical and gene expression differences were only due to biological and technical noise. We repeatedly sampled 10 or 29 cells, respectively, from the pool of cells, which did not express *pdgfra* or *cd24* and counted for each gene the frequency of samples which had equal or higher gene expression compared to the two subpopulations. To account for multiple hypothesis testing we used the Benjamini-Hochberg procedure and set the maximal false discovery rate to 0.05. Additionally, we required a minimal fold-change of 2 for a gene to be accepted as differentially over-expressed. Finally, we considered only genes which were defined as transcriptional regulators by gene ontology (GO) term annotation (GO:0003700, GO:0044212, GO:0045944, GO:0006355, GO: 0000981).

We combined the transcriptional regulators identified in this way with pluripotency network factors ^30^ to arrive at a set of transcription factors which are likely relevant for the lineage decision studied here. For the calculation of co-expression (Fig. 4b-c) we considered a gene to be expressed at normalized RPKM values over 1.

### Single-molecule FISH

Cells growing in gelatinized tissue culture dishes were washed twice with PBS, detached with Accutase (Life technologies) and resuspended in full medium. Formaldehyde was added to the cell suspension to a final concentration of 4%. Cells were incubated for 12 min at room temperature while being rotated and then spun down for 3 min at 1,000rpm. Subsequently cells were permeabilized at least over night in 70% ethanol. For hybridization and imaging cells were attached to chambered cover slides (Nunc Lab-Tek) coated with poly-l-lysine.In the case of intact colonies, adherent cells were fixed for 15 min with 4% formaldehyde by adding formaldehyde to the growth medium and subsequently permeabilized in 70% ethanol.

Oligonucleotide libraries with 20-nt probes for *nanog*, *sox2, ki67, gbx2, tbx3, gata6* and *pax6* were designed and fluorescently labeled as previously described^26^. The hybridization buffer used for smFISH contained 2 × SSC buffer, 25% or 40% formamide, 10% Dextran Sulphate (Sigma), E. coli tRNA (Sigma), Bovine Serum Albumin (Ambion) and Ribonucleoside Vanadyl Complex (New England biolabs). 50 ng - 75 ng of the desired probes were used per 100 μl of hybridization buffer. (The mass refers only to pooled oligonucleotides, excluding fluorophores, and is based on absorbance measurements at 260 nm). Probes were hybridized for 16 - 18 h at 30°C, after which we washed cells twice for 30 min at 30°C in wash buffer (2 × SSC, 25% formamide (for all probes except *gbx2* and *tbx3*) or 40% formamide (for *gbx2* and *tbx3*)), supplemented with Hoechst 33342. For microscopy, we filled the hybridization chamber with a mounting solution containing 1 x PBS, 0.4% Glucose, 100 μg/ml Catalase, 37 μg/ml Glucose Oxidase, and 2 mM Trolox. Imaging was done exactly as described previously ^40^ and home-made MATLAB scripts were used for image analysis. Cells positive for one of the assayed genes were classified as shown in Supplementary Fig.7b.

### Quantification of the flow cytometry experiments

The distribution of cells in the space of *cd24* and *pdgfra* expression was modeled by the sum of 4 bivariate normal distributions. This model has in principle 19 free parameters (8 for the means, 8 for the standard deviations and 3 for the size of the relative contributions). To ensure robust fitting to the date we reduced the number of parameters to 9 by keeping the standard deviations constant and only allowing 4 different values for the means.

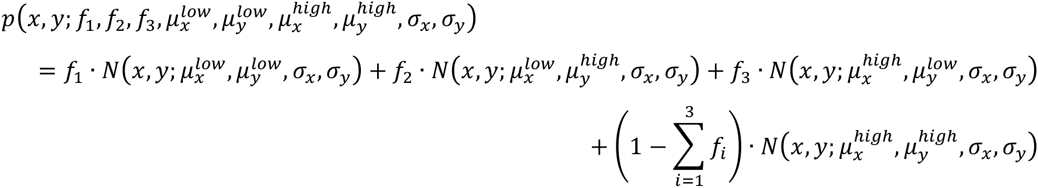

N(x,y,μx,μy,σx,σy) is a bivariate normal distribution in x and y (*pdgfra* and *cd24* expression, respectively) with mean (μ_x_,μ_y_) and standard deviation (σ_x_,σ_y_). This model was fit to a reference data set (typically untreated control cells after 96 h of RA exposure) by maximizing the log-likelihood −log(p). To subsequently calculate the size of the fractions f_i_ for a particular sample we first calculated the probabilities that the expression values (x,y) found in a particular cell were drawn from one the 4 normal distributions N(x,y,μ_x_,μ_y_,σ_x_,σ_y_). The cell was then ascribed to the distribution from which it was most likely drawn.

### Stochastic simulation of the bifurcation

We simulated the bifurcation process using a discretized version of the Langevin equation describing the system (Euler method):

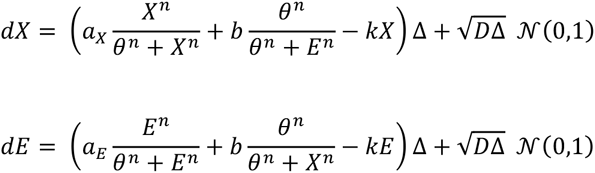

X and E indicate the expression levels of the XEN and ectoderm programs respectively. N(0,1) indicates a Wiener process with mean 0 and standard deviation 1. D sets the strength of gene expression noise and Δ determines the size of the time step. After initializing X and E randomly between 0 and 0.1 we first equilibrated the system for 100 iterations. Subsequently, we propagated the system for 200 additional iterations. To relate the simulation to experimental time scales, the end point of the simulation was taken to be at 96 h. To model the exit from pluripotency the degradation parameter k was switched from a high value (k=10) to a low value (k=1) after 12 h (25 iterations), which allowed X and E to increase. To model timed application of RA the auto-activation parameter for the XEN program aX was switched at various points in time (no RA: a_X_ = 0_;;_ RA: a_X_ = 0.4). For each condition we generated 10000 trajectories and counted the number of trajectories that ended at the XEN or ectoderm attractor (see Supplementary Fig. 5b). The relative frequency of trajectories ending at the XEN attractor is reported in Fig. 3e.

### Used parameters

n = 4

θ = 0.5

Δ = 0.05

pluripotency: k = 10

differentiation: k = 1

a_E_ = 0.5

no RA:, a_X_ = 0

RA: a_X_ = 0.4

low noise: D = 0.01

high noise: D = 0.1

## ACKNOWLEDGEMENTS

SS was supported by the Netherlands Organisation for Scientific Research (NWO/OCW), as part of the Frontiers of Nanoscience program and by an NWO Rubicon award. JG was supported by the Boehringer Ingelheim Foundation as well as a Jerome and Florence Brill Graduate Student Fellowship. RJ was supported by NIH grants HD 045022 and RO1-CA084198. AvO was supported by an ERC Advanced grant (ERC-AdG 294325-GeneNoiseControl) and an NWO Vici award.

---

**Supplementary figure 1.**
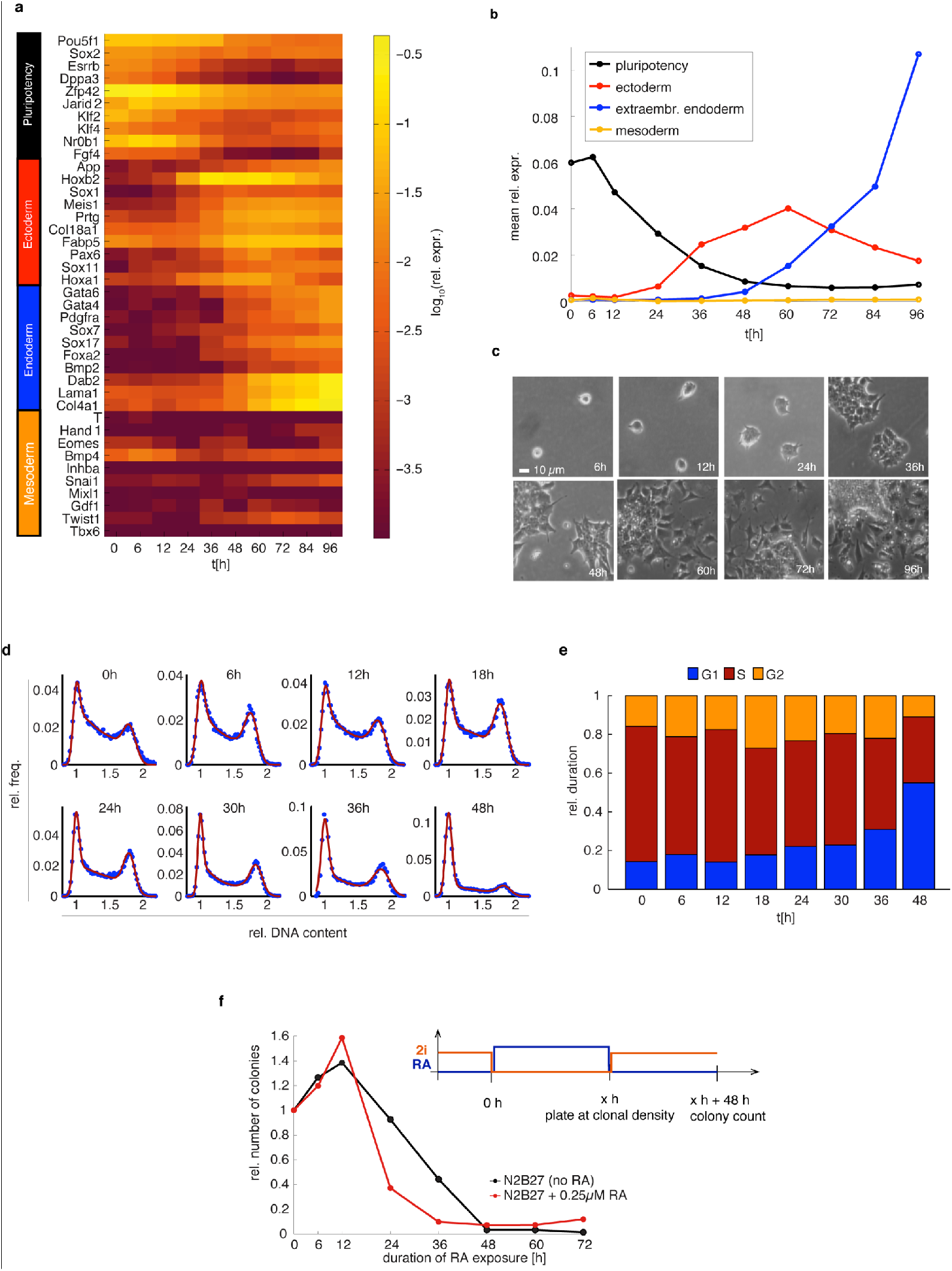
Bulk RNA-seq and phenotypic assays showed that mESCs exited from pluripotency after 12 h of RA exposure. **a**, Gene expression of marker genes for pluripotency and the three germ layers measured by bulk RNA-seq. Expression was normalized to *gapdh* expression at each time point. **b**, Average expression profiles (relative to *gapdh*) of marker genes shown in **a. c**, Cell morphology throughout the differentiation time course. Cells were continuously exposed to 0.25 μM RA. Shown are representative phase contrast images. **d**, Histograms of the relative DNA content measured by Hoechst 33342 staining and flow cytometry (blue symbols) after the indicated periods of exposure to RA. The red solid lines are fits of the Dean-Jet-Fox model ^34^ to the data for individual time points. **e**, Relative lengths of cell cycle phases throughout the time course as determined from fits of the Dean-Jet-Fox model to the measured DNA content. **f**, Clonogenicity after exposure to RA (red data points) or N2B27 basal medium (black data points). After differentiation for the indicated amounts of time cells were replated at a defined, clonal density in 2i medium. After culture for two additional days in 2i medium colonies were counted. The reported values are relative to the number of colonies obtained without differentiation prior to culture in 2i. The inset shows a schematic of this experiment.

**Supplementary figure 2.**
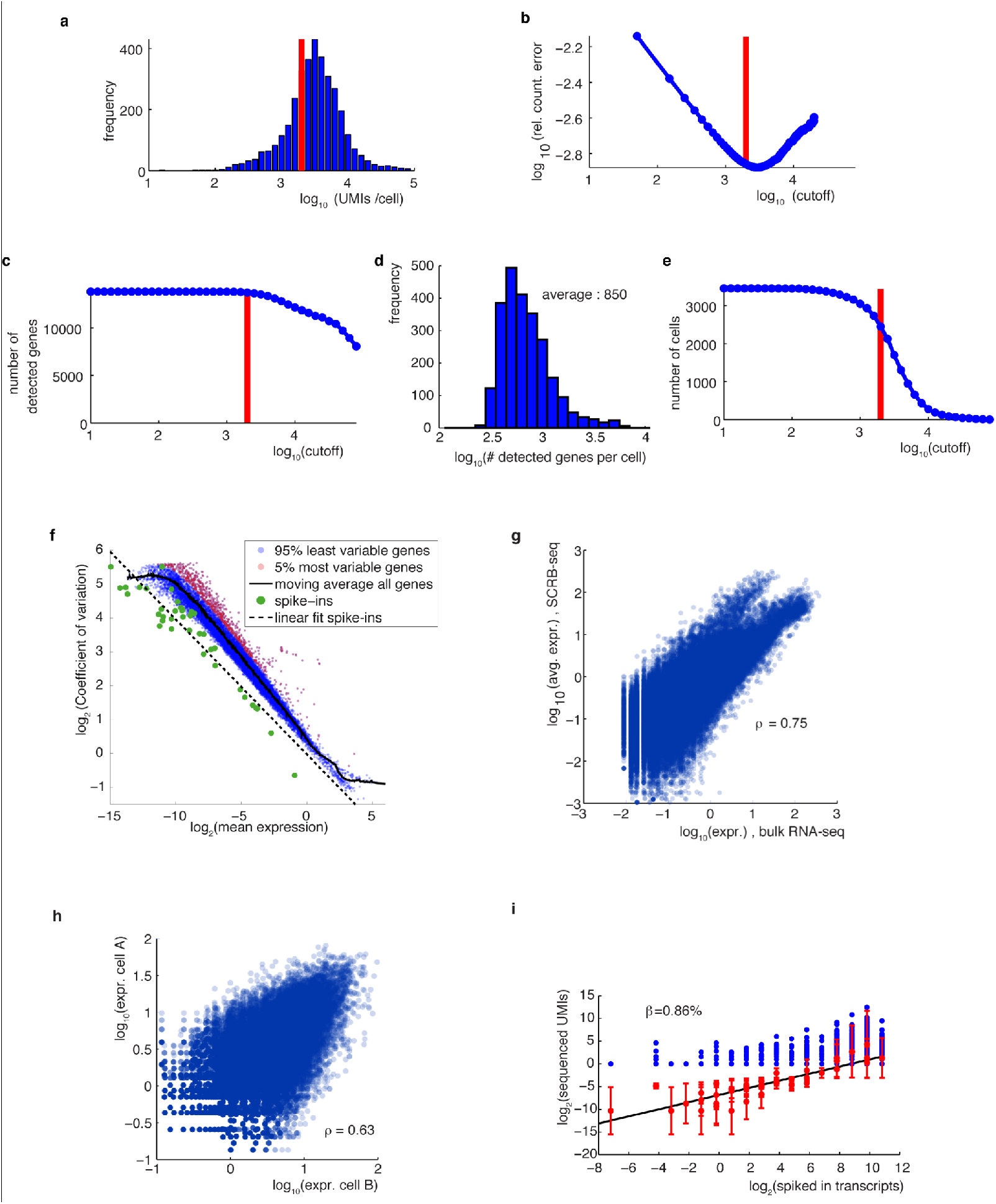
Characterization of the SCRB-seq method. **a**, Frequency of total number of UMIs detected per cell. The red line indicates the cutoff (total UMIs > 2000) above which cells were used for further analysis. **b**, Estimated upper bound for the relative counting error of UMIs with respect to the UMI cutoff. **c**, Number of genes detected across all cells (UMI > 1 in more than 1 cells) with respect to total UMI cutoff. **d**, Distribution of number of detected genes (UMI > 1) per cell for cells with at least 2000 UMIs in total. The average is 850. **e**, Number of cells used for further analysis with respect to total UMI cutoff **f**, Coefficient of variation (CV) of individual genes with respect to mean expression level across all time points. The solid line is a moving average. Indicated in red are the genes, which are considered the 5% most variable taking into consideration the general trend. For comparison, the CVs of spiked in ERCC transcripts are shown in green and a linear fit to these data points is shown as a dashed, black line. **g**, Comparison of bulk RNA-seq measurements and SCRB-seq measurements averaged over cells for individual time points. Pearson correlation ρ = 0.75. **h**, Comparison of expression levels measured by SCRB-seq in 100 randomly selected pairs of single cells in 2i conditions. Pearson correlation ρ = 0.63. **i**, Number of spiked in ERCC transcripts with respect to sequenced spike-in UMIs. The blue symbols show the measurements while red symbols indicate the mean; whiskers indicate standard deviations. From a linear fit of the data the recovery efficiency is determined to be 0.9%. The slope of the linear fit is 0.78.

**Supplementary figure 3.**
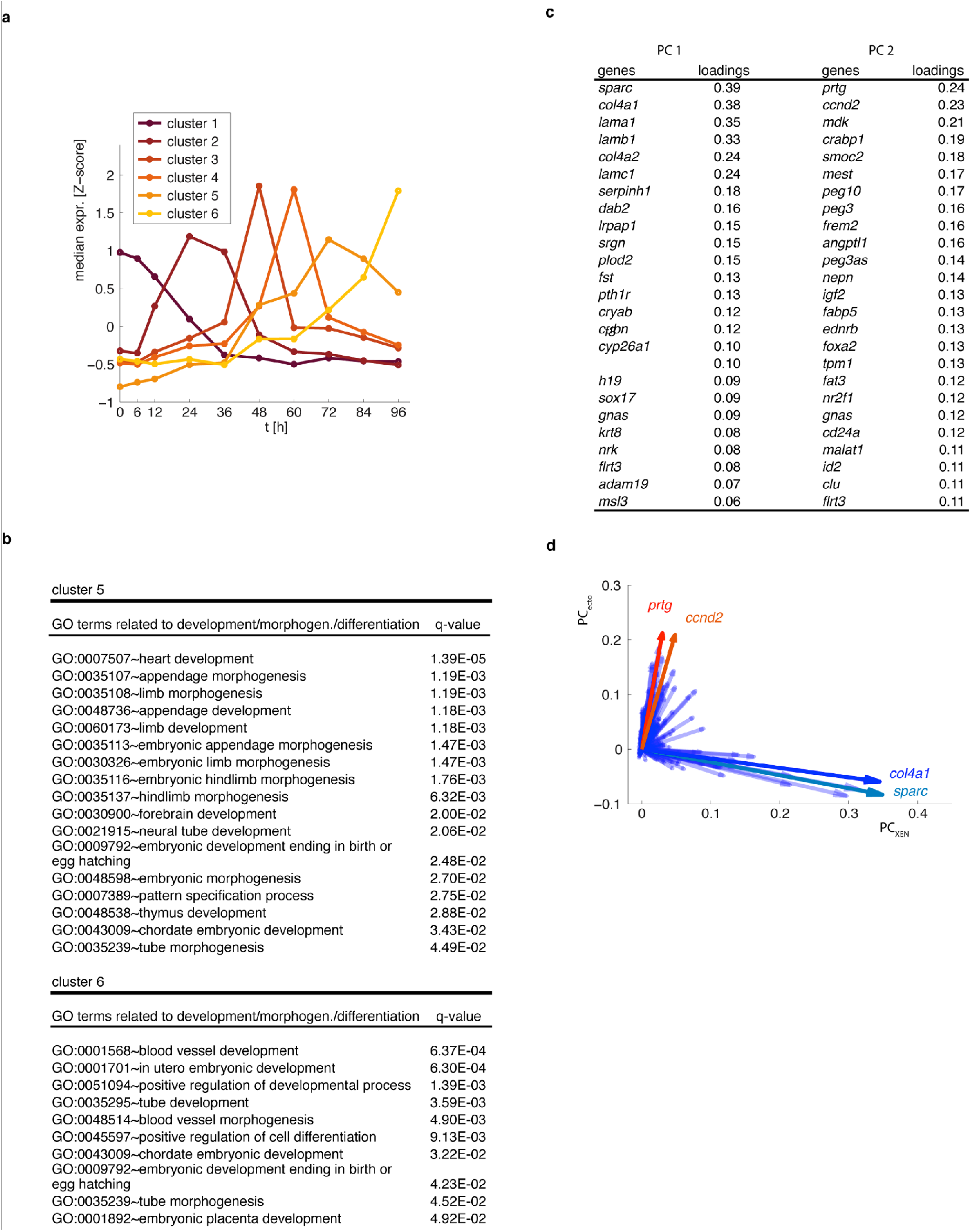
Principal component analysis of the differentiation dynamics. **a**, Expression profiles measured by bulk RNA-seq were clustered by k-means clustering and 6 robust clusters were identified. The median of the expression profiles in each cluster is shown. Clusters were ordered by the position of the expression maximum. **b**, GO terms which were significantly enriched in clusters 5 and 6 (q-value < 5%). **c**, Loadings of the 25 genes with the highest loadings in the first two principal components (PC 1 and PC 2). **d**, Graphical representation of the data in **e**. Each arrow represents the loading of a gene. The elements of the vector defining the arrow are the gene’s loadings.

**Supplementary figure 4.**
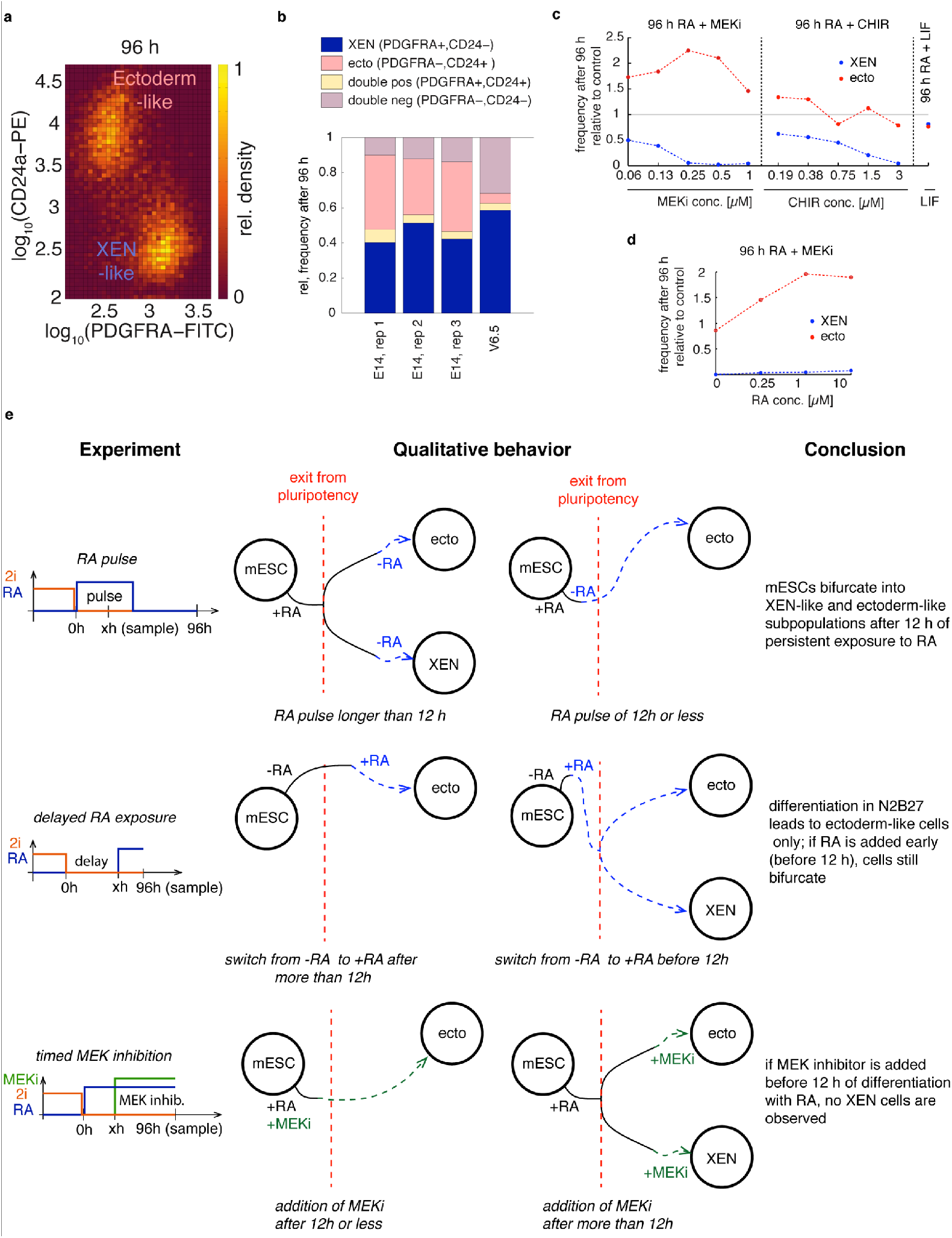
Cell type quantification by antibody staining and timed lineage cue experiments. **a**, Flow cytometry measurement of cells stained with *cd24* and *pdgfra* antibodies after 96 h of RA exposure. The heatmap represents the relative density of cells. **b**, Frequency of XEN-like and ectoderm-like cells after 96 h exposure to RA. Cells were classified based on *cd24* and *pdgfra* expression measured by antibody staining and flow cytometry. Shown are results for 3 biological replicates with E14 mESCs as well as V6.5 mESCs. **c**, Frequency of XEN-like and ectoderm-like cells after 96 h exposure to RA and MEK inhibitor PD0325901 (MEKi), RA and GSK3 inhibitor CHIR99021 (CHIR) or RA and LIF relative to the control (RA only). Cells were classified based on *cd24* and *pdgfra* expression measured by antibody staining and flow cytometry. **d**, Frequency of XEN-like and ectoderm-like cells after 96 h exposure to various concentrations of RA and 0.5 μM MEK inhibitor PD0325901 (MEKi) relative to the control (0.25 μM RA only). Cells were classified based on *cd24* and *pdgfra* expression measured by antibody staining and flow cytometry. **e,** Schematic overview of experiments and qualitative results. ‘mESC’ : pluripotent cells growing in 2i medium, ‘ecto’ : ectoderm-like cells, ‘XEN’ : XEN-like cells, ‘jam’ : undifferentiated jammed cells, ‘MEKi : MEK inhibitor, ‘RA’ : retinoic acid, ‘2i’ : 2i medium

**Supplementary figure 5.**
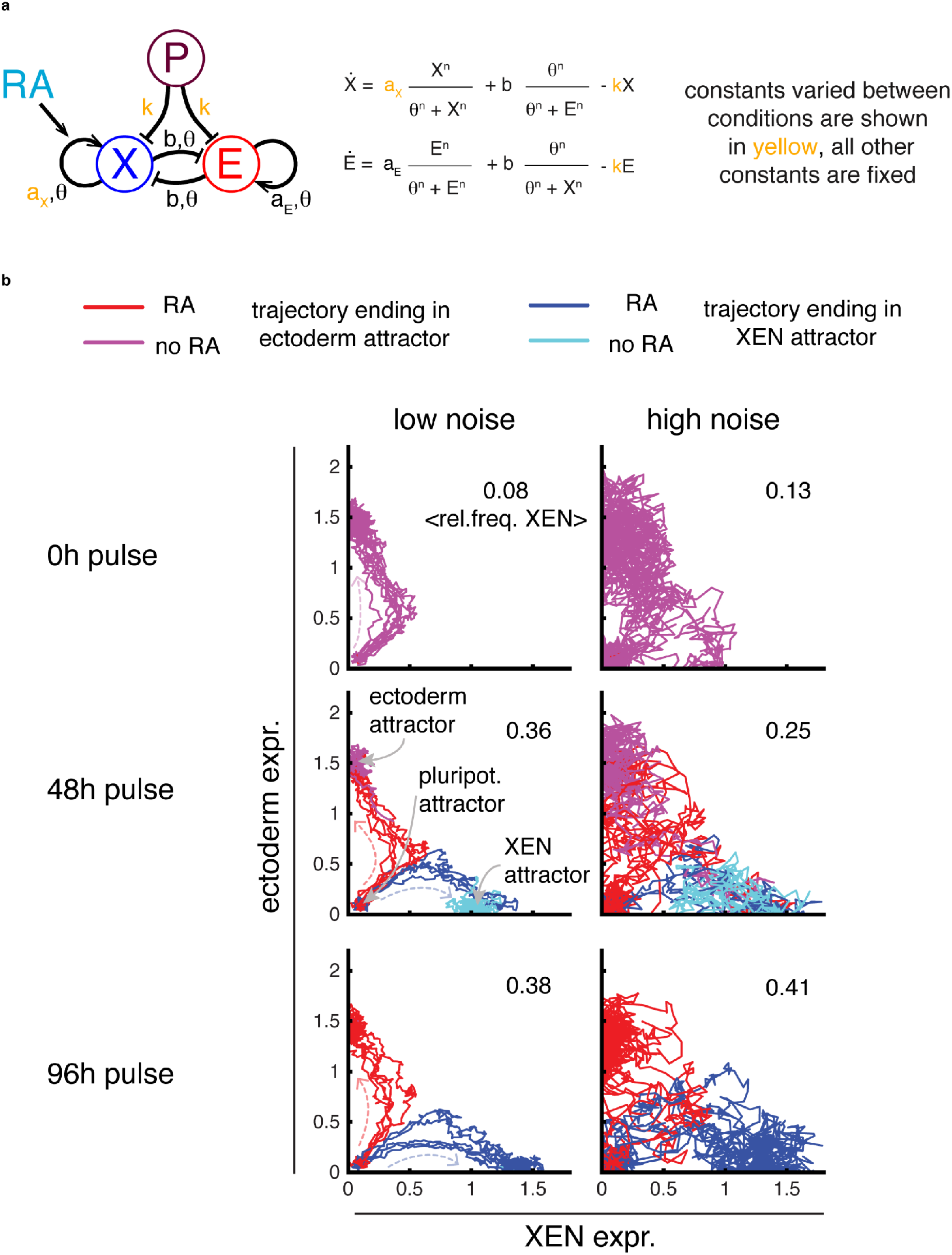
Stochastic simulation of the bifurcation process. **a**, Schematic representation and equations defining a minimal gene regulatory network that can produce a lineage bifurcation ^20^. Pointy arrows indicate (auto)activation; blunted arrows indicate repression. E and X represent expression of ectoderm-like and XEN-like transcriptional programs, respectively. P stands for the pluripotency network. Parameters which are changed between conditions are shown in yellow. All other parameters are fixed. **b**, Exemplary trajectories for the case of an RA pulse. Each panel shows 10 trajectories simulated under the condition indicated at each row and column. All trajectories start at the pluripotency attractor and end either at the XEN or ectoderm attractor. Dashed arrows indicate their overall direction. Trajectories are colored according to their endpoint and the presence of RA. Red-violet trajectories end at the ectoderm attractor, cyan-blue trajectories end at the XEN attractor. During the red and blue parts of trajectories RA is present, during the violet and cyan parts of the trajectories there is no RA. The number in each panel gives the relative frequency of trajectories ending at the XEN attractor, where 10000 trajectories were simulated.

**Supplementary figure 6.**
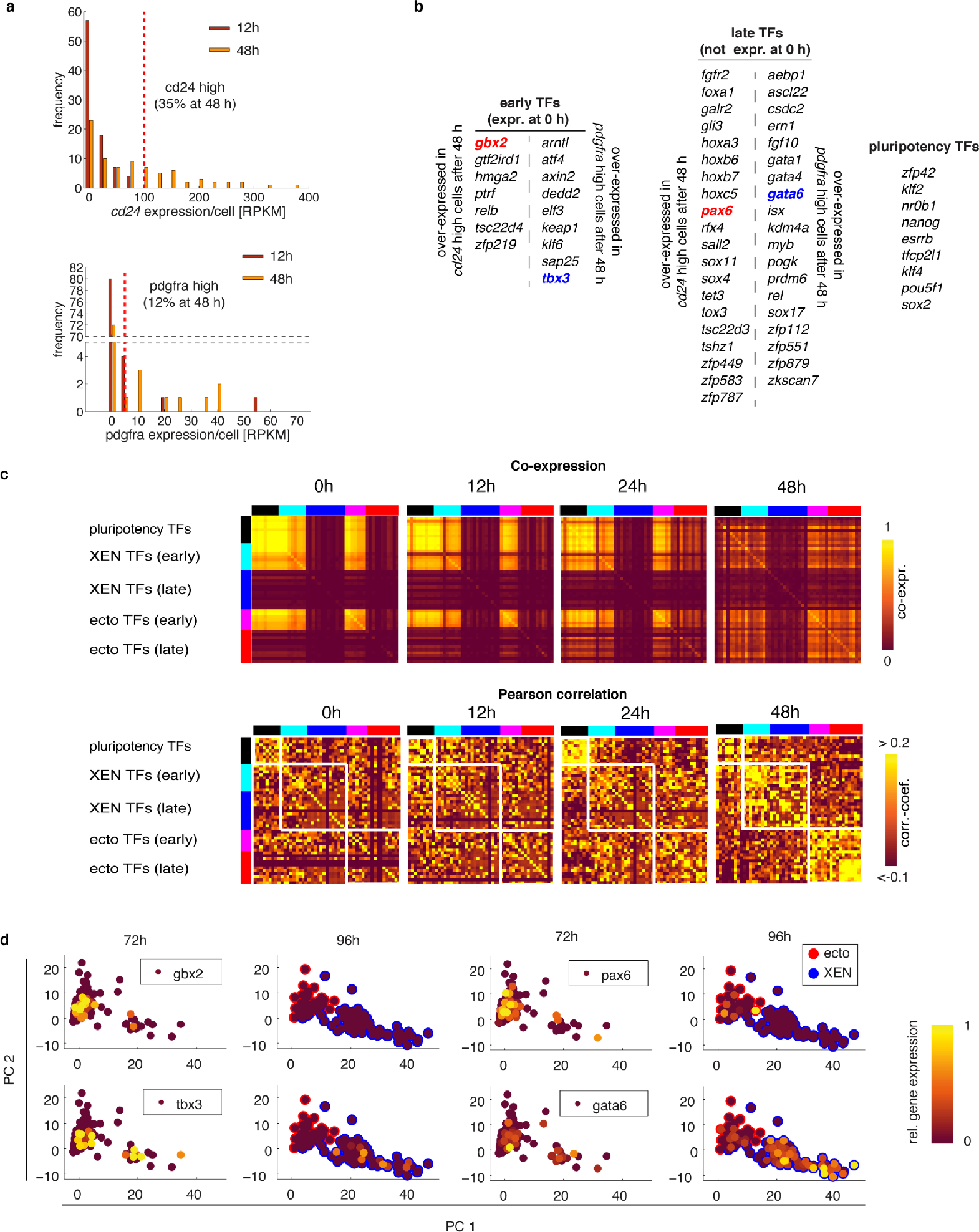
Identification of differentially expressed transcription factors with SMART-seq. **a**, Single-cell expression of *cd24* and *pdgfra* after 12 h and 48 h exposure to RA, measured by SMART-seq. The red dashed line indicates the expression level that defines *cd24* or *pdgfra* high cells. **b**, List of transcriptional regulators over-expressed after 48 h of RA exposure in *pdgfra* or *cd24* high cells. The transcription factors shown in color were picked for further analysis. **c**, Co-expression and Pearson correlation between a set of core transcriptional regulators after 0 h, 12 h, 24 h and 48 h of RA exposure. The gene set comprised the differentially expressed transcriptional regulators identified here (see Fig. 4 a), as well as pluripotency related transcription factors listed in ^30^ (see **b**). Co-expression and Pearson correlation were calculated using gene expression measured by SMART-seq. Co-expression of two genes was quantified as the fraction of cells with significant expression of both genes. Correlation coefficients exceeding the range [−0.1,0.2] are shown in the same color as the respective extreme values of that range. **d**, Single-cell gene expression of *pax6*, *gata6*, *gbx2* and *tbx3* after 72 h and 96 h of RA exposure, measured with SCRB-seq. The position of each data point represents the expression profile of a cell in the space of the first two principal components (PC^XEN^ and PC_ecto_). The color of a data point reflects the expression of the indicated gene, relative to the maximal expression across all cells and the two time points. Cells belonging to the ectoderm-like and an XEN-like cluster after 96 h of RA exposure are indicated by red or blue edges, respectively.

**Supplementary figure 7.**
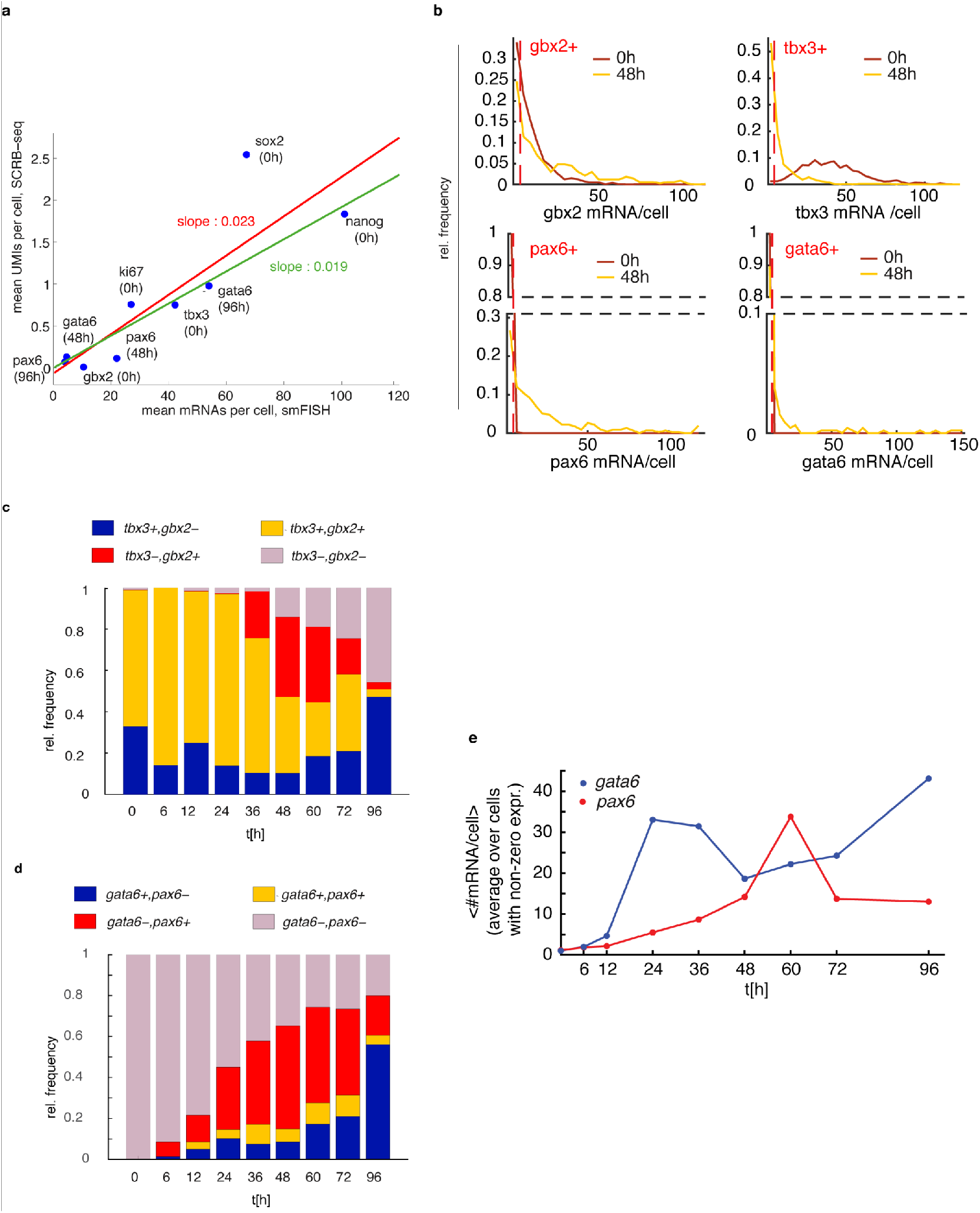
Expression dynamics of transcription factors measured by smFISH. **a**, Sensitivity of smFISH compared to SCRB-seq. Each data point is the single-cell expression level of the indicated gene (at the indicated time point) averaged over all measured cells. The solid lines are linear fits, where the red model had an arbitrary intercept and the green model was forced to go through the origin. smFISH detected approximately 40 times more transcripts than SCRB-seq. **b**, Abundances of *pax6*, *gata6*, *gbx2* and *tbx3* mRNAs after 0 h and 48 h of RA exposure measured with smFISH. The red dashed line indicates the expression levels that demarcates cells, which are considered positive (“+”) for the indicated gene. **c**, Fractions of cells which (co)expressed *gbx2* and *tbx3* throughout the differentiation time course, based on the smFISH measurements shown in Fig. 5c Thresholds for significant expression are indicated in Supplementary Fig. 9b. **d**, Fractions of cells which (co)expressed *pax6* and *gata6* throughout the differentiation time course, based on the smFISH measurements shown in Fig. 5d. Thresholds for significant expression are indicated in Supplementary Fig. 9b. **e**, Number of *pax6* or *gata6* transcripts per cell averaged over cells that have at least one *pax6* or one *gata6* transcript, respectively. Transcript abundance was measured by smFISH. Cells were exposed to RA acid for the indicated periods of time.

## REFERENCES

1. Cohen, D. E. & Melton, D. Turning straw into gold: directing cell fate for regenerative medicine. Nat Rev Genet 12 243–252 (2011).

2. Soldner, F. & Jaenisch, R. iPSC Disease Modeling. Science 338 1155–1156 (2012).

3. Tabar, V. & Studer, L. Pluripotent stem cells in regenerative medicine: challenges and recent progress. Nat Rev Genet 15 82–92 (2014).

4. Semrau, S. & van Oudenaarden, A. Studying Lineage Decision-Making In Vitro: Emerging Concepts and Novel Tools. Annu. Rev. Cell Dev. Biol. 31 317–345 (2015).

5. Balázsi, G., van Oudenaarden, A. & Collins, J. J. Cellular Decision Making and Biological Noise: From Microbes to Mammals. Cell 144 910–925 (2011).

6. Pauklin, S. & Vallier, L. The Cell-Cycle State of Stem Cells Determines Cell Fate Propensity. Cell 155 135–147 (2013).

7. Kobayashi, T. et al. The cyclic gene Hes1 contributes to diverse differentiation responses of embryonic stem cells. Gene Dev 23 1870–1875 (2009).

8. Morgani, S. M. & Brickman, J. M. The molecular underpinnings of totipotency. Philos. Trans. R. Soc. Lond., B, Biol. Sci. 369 20130549–20130549 (2014).

9. Ying, Q.-L. et al. The ground state of embryonic stem cell self-renewal. Nature 453 519–523 (2008).

10. Capo-chichi, C. D. et al. Perception of differentiation cues by GATA factors in primitive endoderm lineage determination of mouse embryonic stem cells. Developmental biology 286 574–586 (2005).

11. Rhinn, M. & Dolle, P. Retinoic acid signalling during development. Development 139, 843–858 (2012).

12. Soumillon, M., Cacchiarelli, D. & Semrau, S. Characterization of directed differentiation by high-throughput single-cell RNA-Seq. bioRxiv (2014).

13. Marks, H. et al. The Transcriptional and Epigenomic Foundations of Ground State Pluripotency. Cell Stem Cell 149 590–604 (2012).

14. Hennig, C. Cluster-wise assessment of cluster stability. Computational Statistics & Data Analysis 52 258–271 (2007).

15. Johnson, A. T. et al. Synthesis and Characterization of a Highly Potent and Effective Antagonist of Retinoic Acid Receptors. J. Med. Chem. 38 4764–4767 (1995).

16. Pruszak, J., Ludwig, W., Blak, A., Alavian, K. & Isacson, O. CD15, CD24 and CD29 Define a Surface Biomarker Code for Neural Lineage Differentiation of Stem Cells. STEM CELLS N/A–N/A (2009). doi:10.1002/stem.211

17. Artus, J., Panthier, J.-J. & Hadjantonakis, A.-K. A role for PDGF signaling in expansion of the extra-embryonic endoderm lineage of the mouse blastocyst. Development 137 3361–3372 (2010).

18. Hamilton, W. B. & Brickman, J. M. Erk signaling suppresses embryonic stem cell self-renewal to specify endoderm. Cell Reports 9 2056–2070 (2014).

19. Schröter, C., Rué, P., Mackenzie, J. P. & Arias, A. M. FGF/MAPK signaling sets the switching threshold of a bistable circuit controlling cell fate decisions in embryonic stem cells. Development 142 4205–4216 (2015).

20. Huang, S., Guo, Y.-P., May, G. & Enver, T. Bifurcation dynamics in lineage-commitment in bipotent progenitor cells. Developmental biology 305 695–713 (2007).

21. Picelli, S. et al. Smart-seq2 for sensitive full-length transcriptome profiling in single cells. Nat Methods 10 1096–1098 (2013).

22. Liu, A. & Joyner, A. L. EN and GBX2 play essential roles downstream of FGF8 in patterning the mouse mid/hindbrain region. Development 128 181–191 (2001).

23. Osumi, N., Shinohara, H., Numayama-Tsuruta, K. & Maekawa, M. Concise Review: Pax6 Transcription Factor Contributes to both Embryonic and Adult Neurogenesis as a Multifunctional Regulator. STEM CELLS 26 1663–1672 (2008).

24. Lu, R., Yang, A. & Jin, Y. Dual Functions of T-Box 3 (Tbx3) in the Control of Self-renewal and Extraembryonic Endoderm Differentiation in Mouse Embryonic Stem Cells. J Biol Chem 286 8425–8436 (2011).

25. Arnold, S. J. & Robertson, E. J. Making a commitment: cell lineage allocation and axis patterning in the early mouse embryo. Nature Reviews Molecular Cell Biology 10 91–103 (2009).

26. Raj, A., van den Bogaard, P., Rifkin, S. A., van Oudenaarden, A. & Tyagi, S. Imaging individual mRNA molecules using multiple singly labeled probes. Nat Methods 5 877–879 (2008).

27. Klein, A. M. et al. Droplet Barcoding for Single-Cell Transcriptomics Applied to Embryonic Stem Cells. Cell 161 1187–1201 (2015).

28. Turner, D. A., Trott, J., Hayward, P., Rue, P. & Martinez Arias, A. An interplay between extracellular signalling and the dynamics of the exit from pluripotency drives cell fate decisions in mouse ES cells. Biology Open 3 614–626 (2014).

29. Marco, E. et al. Bifurcation analysis of single-cell gene expression data reveals epigenetic landscape. Proceedings of the National Academy of Sciences 111, E5643–E5650 (2014).

30. Dunn, S. J., Martello, G., Yordanov, B., Emmott, S. & Smith, A. G. Defining an essential transcription factor program for naive pluripotency. Science 344 1156–1160 (2014).

31. Thomson, M. et al. Pluripotency Factors in Embryonic Stem Cells Regulate Differentiation into Germ Layers. Cell 145 875–889 (2011).

32. Trott, J. & Martinez Arias, A. Single cell lineage analysis of mouse embryonic stem cells at the exit from pluripotency. Biology Open 2 1049–1056 (2013).

33. Singer, Z. S. et al. Dynamic Heterogeneity and DNA Methylation in Embryonic Stem Cells. Molecular cell 55 319–331 (2014).

34. Fox, M. H. A model for the computer analysis of synchronous DNA distributions obtained by flow cytometry. Cytometry 1 71–77 (1980).

35. Picelli, S. et al. Full-length RNA-seq from single cells using Smart-seq2. Nature Protocols 9 171–181 (2014).

36. Trombetta, J. J. et al. Preparation of Single-Cell RNA-Seq Libraries for Next Generation Sequencing. Curr Protoc Mol Biol 107 4.22.1–4.22.17 (2014).

37. Trapnell, C. et al. Differential gene and transcript expression analysis of RNA-seq experiments with TopHat and Cufflinks. Nature Protocols 7 562–578 (2012).

38. Huang, D. W., Sherman, B. T. & Lempicki, R. A. Systematic and integrative analysis of large gene lists using DAVID bioinformatics resources. Nature Protocols 4 44–57 (2008).

39. van der Maaten, L. & Hinton, G. Visualizing Data using t-SNE. Journal of Machine Learning Research 9 2579–2605 (2008).

40. Semrau, S. et al. FuseFISH: Robust Detection of Transcribed Gene Fusions in Single Cells. Cell Reports 6 18–23 (2014).

